# A parabrachial hub for the prioritization of survival behavior

**DOI:** 10.1101/2024.02.26.582069

**Authors:** Nitsan Goldstein, Amadeus Maes, Heather N. Allen, Tyler S. Nelson, Kayla A. Kruger, Morgan Kindel, Nicholas K. Smith, Jamie R.E. Carty, Rachael E. Villari, Ella Cho, Erin L. Marble, Rajesh Khanna, Bradley K. Taylor, Ann Kennedy, J. Nicholas Betley

## Abstract

Long-term sustained pain in the absence of acute physical injury is a prominent feature of chronic pain conditions. While neurons responding to noxious stimuli have been identified, understanding the signals that persist without ongoing painful stimuli remains a challenge. Using an ethological approach based on the prioritization of adaptive survival behaviors, we determined that neuropeptide Y (NPY) signaling from multiple sources converges on parabrachial neurons expressing the NPY Y1 receptor to reduce sustained pain responses. Neural activity recordings and computational modeling demonstrate that activity in Y1R parabrachial neurons is elevated following injury, predicts functional coping behavior, and is inhibited by competing survival needs. Taken together, our findings suggest that parabrachial Y1 receptor-expressing neurons are a critical hub for endogenous analgesic pathways that suppress sustained pain states.

## Main Text

Despite progress in understanding neural pathways that process the sensory and emotional aspects of pain (*1, 2*), the neural circuits engaged during long-term pain states remain elusive (*3*). Populations of neurons that are activated during painful stimuli have been identified throughout the peripheral and central nervous systems (*4–11*). Furthermore, altered spinal circuitry following injury is well documented and, at least in part, underlies transitions to chronic pain (*12–14*). It is likely that potentiated spinal pathways converge on a central neural node that signals the multimodal sensory and affective components of long-term pain. Identification of neurons that are tuned to pain state, and not just responsive to transient noxious stimuli, could serve as a gateway to understand how changes in neural activity lead to maladaptive, prolonged pain (*15, 16*).

How can we find state-tuned neurons in the brain? Consistent changes in neural activity during persistent pain are not clearly discernible from human imaging studies (*3*). One approach is to uncover situations in which the global pain state can be altered and identify the neurons and circuits underlying these changes (*17–19*). Hunger suppresses behavioral responses and the negative affective state that follow tissue damage (*19*), suggesting that competing need states may activate endogenous analgesic circuits to prioritize survival behaviors. Understanding the mechanisms used to suppress pain during competing needs could allow for the identification of neuron populations that encode a sustained pain state. Here, we show that a variety of ethologically relevant need states suppress sustained pain responses. We identified common circuit and molecular mechanisms that led us to a population of parabrachial neurons expressing the neuropeptide Y (NPY) Y1 receptor. Both neural activity monitoring and computational modeling demonstrate that activity in these neurons correlates with lasting pain and is modulated by NPY to suppress long term pain states when animals are faced with competing needs.

### Physiological and environmental needs suppress pain

To explore the possibility that various survival needs can activate endogenous analgesic circuits to suppress lasting pain, we screened for physiological and environmental factors that inhibit prolonged behavioral responses to injury. We found that hunger, thirst, innate fear, and conditioned fear are capable of suppressing behavioral responses to multiple modalities of sustained pain (Fig. 1).

**Figure 1.**
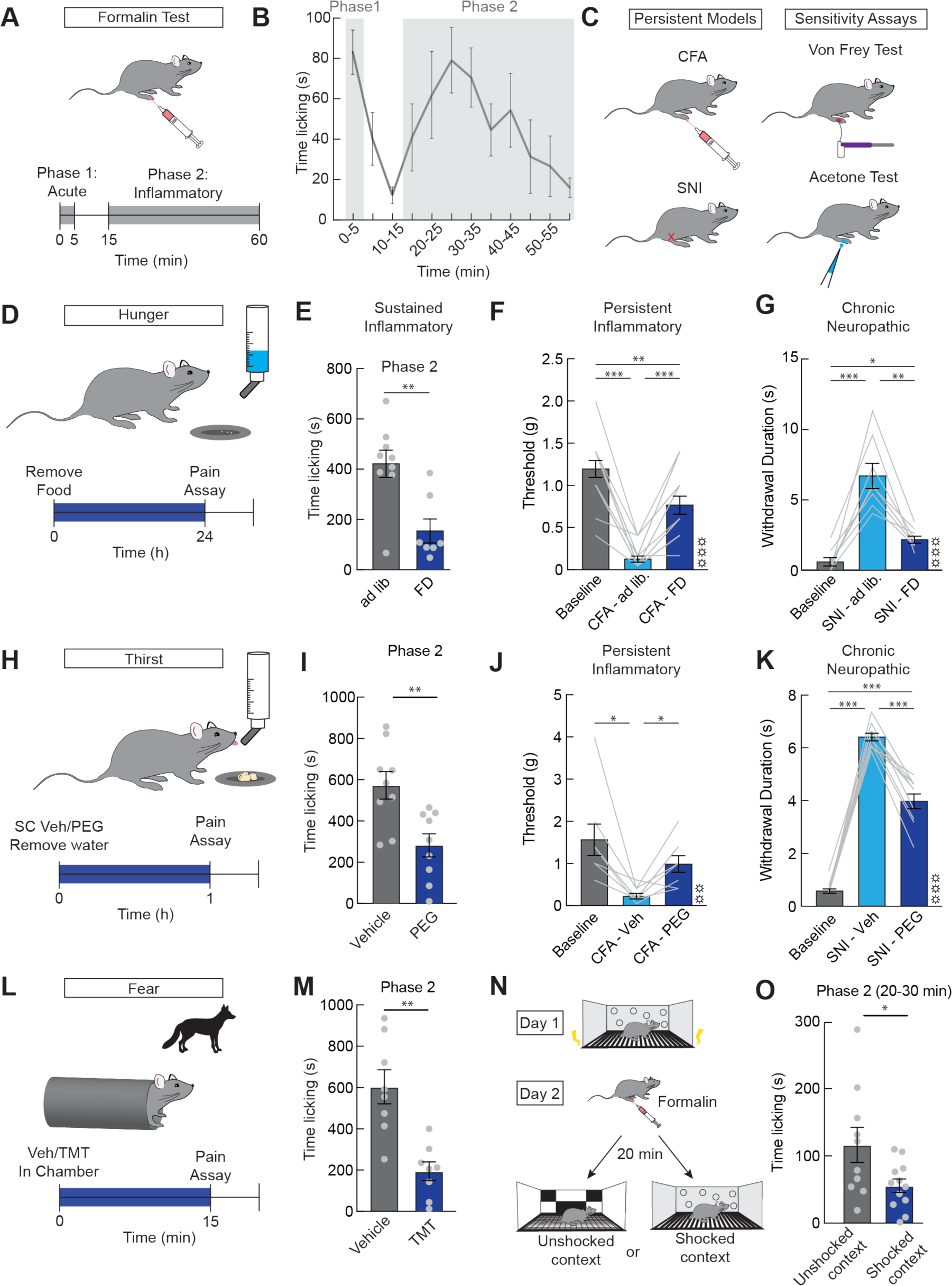
Prolonged pain responses are suppressed by competing needs. (**A**) Formalin was injected into the dorsal surface of the hindpaw, resulting in an initial acute phase (Phase 1, 0-5 min) and a sustained phase (Phase 2, 15-60 min). (**B**) Time spent licking paw in 5 min bins after an injection of formalin (n=9) showing initial and sustained phases. (**C**) Complete Freund’s Adjuvant (CFA) was used as a model of persistent inflammatory pain and spared nerve injury (SNI) was used as a model of neuropathic pain. The von Frey test was used to assess mechanical allodynia in both models and the acetone test was used to test cold allodynia after SNI. (**D**) Timeline for 24 h food deprivation experiments. (**E**) Time licking paw during Phase 2 in *ad libitum* fed mice (grey) and food deprived mice (blue, n=7-9/group, unpaired t-test, p<0.01). (**F**) Withdrawal threshold to mechanical stimuli before CFA (baseline, grey), *ad libitum* fed mice after CFA (light blue), and 24 h food deprived mice after CFA (dark blue, n=15, repeated measures one-way ANOVA, p<0.001). (**G**) Withdrawal duration after acetone application before SNI (baseline, grey), *ad libitum* fed mice after SNI (light blue), and 24 h food deprived mice after SNI (dark blue, n=8, repeated measures one-way ANOVA, p<0.001). (**H**) Timeline for polyethylene glycol (PEG)-induced thirst (30%, 20ul/g body weight s.c.) experiments. (**I**) Time licking paw during Phase 2 in saline treated mice (grey) and PEG treated mice (blue, n=9/group, unpaired t-test, p<0.01). (**J**) Withdrawal threshold to mechanical stimuli before CFA (baseline, grey), control mice treated with saline after CFA (light blue), and mice treated with PEG after CFA (dark blue, n=8, repeated measures one-way ANOVA, p<0.01). (**K**) Withdrawal duration after acetone application before SNI (baseline, grey), control mice treated with saline after SNI (light blue), and mice treated with PEG after SNI (dark blue, n=10, repeated measures one-way ANOVA, p<0.001). (**L**) Timeline for experiments during innate fear (exposure to Trimethylthiazoline (TMT, fox odor)). (**M**) Time licking paw during Phase 2 in mice with control PBS in the chamber (grey) and TMT in the chamber (blue, n=8/group, unpaired t-test, p<0.01). (**N**) Timeline of experiments during conditioned fear. Conditioning involved pairing a shock with a context on day 1. On day 2, mice were injected with formalin and then placed in either the conditioned or novel context 20 minutes after the injection. Paw licking was scored for 10 min during the inflammatory phase. (**O**) Time licking 20-30 min after formalin injection while animals were in either in the unshocked context (grey) or the shocked context (blue, n=10-12/group, unpaired t-test, p<0.05). Data are expressed as mean ± SEM. Grey dots represent individual mice in between subject experiments and grey lines represent individual mice in within subjects experiments. T-test and post-hoc comparisons: *p<0.05, **p<0.01, ***p<0.001. ANOVA main effect of group: ☼☼p<0.01, ☼☼☼p<0.001

We used a variety of assays to assess nociceptive responses across different timescales, injury sources, and types of allodynia under competing need states (Fig. 1A-C). 24 h food deprivation strongly suppresses wound licking in the inflammatory (second) phase of formalin induced pain (Fig. 1D-E, Fig. S1A), mechanical allodynia due to persistent inflammation (Complete Freund’s Adjuvant, CFA, Fig. 1F, Fig. S1E-F), and mechanical and cold allodynia due to spared nerve injury (SNI, Fig. 1G, Fig. S1H-I). The ability of homeostatic need states to suppress pain is not unique to hunger, as we found that inducing hypovolemic thirst with a peripheral injection of polyethylene glycol (PEG, Fig. S2A-B) also reduces both inflammatory phase licking in the formalin assay (Fig. 1H-I, Fig. S1C) and mechanical and cold allodynia (Fig. 1J-K, Fig. S1G, S1J). Unlike the responses during longer term inflammatory or neuropathic sensitization, hunger and thirst did not affect acute (first phase) responses to formalin induced pain (Fig. S1B, S1D).

External threats similarly suppress sustained responses to injury. Introduction of a predator odor (Trimethylthiazoline, TMT) or an environment previously associated with an aversive shock stimulus both attenuate responses to formalin (Fig. 1L-O, Fig. S3A-D, supplementary text). Though these stimuli elicit behaviors such as freezing and avoidance that could interfere with coping behaviors such as paw licking and withdrawal, we find that withdrawal, licking, and jumping responses to noxious heat were unaffected by predator odor (Fig. S3E-H). Moreover, conditioned fear did not affect responses to formalin when mice were placed in the chamber during the acute phase (Fig. S3C-D) demonstrating that animals are still able to attend to noxious stimuli and perform basic coping behaviors (supplementary text). Taken together, these results point to the existence of ethologically relevant analgesic neural pathways.

### Bulk transmission of NPY in the lPBN reduces sustained, long-lasting pain by acting on the Y1 receptor

Given the similarities in the effects of different competing needs on multiple modalities of sustained pain, we reasoned that a shared mechanism may underly these analgesic effects. Because NPY release in lateral parabrachial nucleus (lPBN) during hunger suppresses inflammatory pain responses (*19*), we reasoned that NPY signaling at an lPBN hub may suppress persistent pain during different survival needs. We find that NPY signaling in the lPBN recapitulates the effects of competing needs on models of persistent inflammation as microinfusions of NPY into the lPBN decreased mechanical hypersensitivity without affecting inflammation (Fig. S4). Blocking NPY signaling with a Y1 receptor (Y1R) antagonist during competing needs occluded the analgesic effects of hunger and fear on inflammatory pain behavior (Fig. 2A-D) (*19*). The effect of blocking NPY signaling is specific to sustained pain behavior as we do not observe changes in acute nociception or mobility (Fig. 2C, Fig. S5A). Further, the antagonist did not change noxious responding in the absence of competing needs (Fig. S5B-D), nor did it influence fear responses in the absence of injury (Fig. S5E-F). These results suggest that the endogenous release of NPY in the lPBN mediates analgesia during acute survival threats.

**Figure 2.**
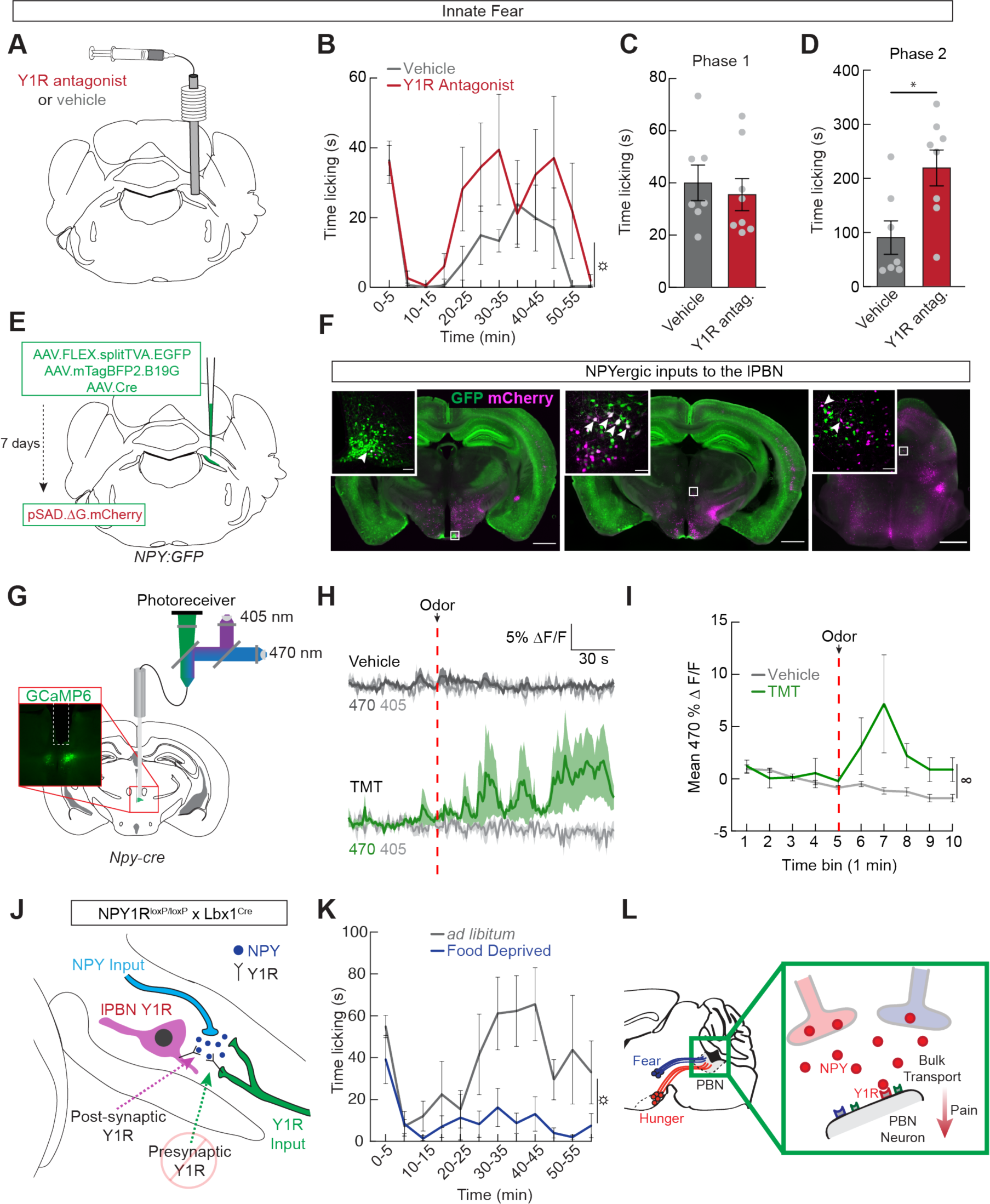
Competing drives suppress pain responses via bulk NPY signaling on lPBN neurons that express the Y1 receptor. (**A**) Microinfusion approach to antagonize NPY-Y1 receptor signaling in the lPBN. (**B**) Time spent licking paw in 5 min bins after an injection of formalin and a control infusion of vehicle (grey) or Y1R antagonist in TMT exposed mice (red, n=7-8/group, repeated measures two-way ANOVA, main effect of group p<0.05). (**C**) Time spent licking paw during Phase 1 in mice exposed to TMT after either a vehicle or Y1R antagonist infusion (n=7-8/group, unpaired t-test, ns). (**D**) Time licking paw during Phase 2 in mice exposed to TMT after either a vehicle or Y1R antagonist infusion (n=7-8/group, unpaired t-test, p<0.05). (**E**) Labeling strategy to identify NPY-expressing neurons projecting to the lPBN. (**F**) Representative images of three regions containing double-labeled NPY-positive (GFP) and rabies-positive (mCherry) neurons that project to the lPBN. Left, arcuate nucleus; middle, subparafascilcular nucleus; right, ventral medial periaqueductal grey. Scale bar, 1 mm (inset, 50 μm). (**G**) Dual wavelength fiber photometry was used to record activity of subparafascilcular (SPF) nucleus NPY-expressing neurons. Inset, representative image showing GCaMP6s expression in SPF NPY neurons below a fiber optic tract. (**H**) Average ΔF/F of GCaMP6s signal from NPY neurons after exposure to control PBS (top, dark grey) or TMT (bottom, green). Signals are aligned to the introduction of odor into the chamber. Dark grey and green, 470 nm; light grey, 405 nm. Dark lines represent mean and lighter, shaded areas represent SEM. (**I**) Mean ΔF/F in 1 min bins of GCaMP6s signal after PBS (grey) or TMT (green) exposure (n=5, repeated measures two-way ANOVA, main effect of group p=0.05, group x time interaction p<0.05). (**J**) Diagram depicting two possible sites of action of NPY in the lPBN: presynaptic inputs from the spinal cord and postsynaptic receptors on lPBN neurons. Y1 receptors were conditionally deleted from the spinal cord and the effect of NPY on pain was tested. (**K**) Time spent licking paw in 5 min bins after an injection of formalin in *ad libitum* fed (grey) and food deprived (blue) spinal cord Y1R conditional knockout mice (n=6, repeated measures two-way ANOVA, main effect of group p<0.05). (**L**) Summary model. Bulk NPY transport acts on Y1 receptors in the lPBN to suppress pain when animals are faced with competing needs. Data are expressed as mean ± SEM. Grey dots represent individual mice. T-test and post-hoc comparisons: *p<0.05. ANOVA main effect of group: ☼p<0.05. ANOVA interaction: ∞p<0.05.

We next sought to identify inputs to the lPBN that could release NPY to suppress pain. We used monosynaptic rabies tracing in NPY-GFP mice to identify NPYergic inputs into the lPBN (Fig. 2E). We identified several populations of NPY neurons that project to the lPBN, including arcuate nucleus NPY-expressing neurons that are active during hunger, as well as previously uncharacterized populations (Fig. 2F). We found that a PBN-projecting NPY population in the subparafascilcular nucleus of the thalamus is activated by innate threat, but not other needs like thirst (TMT, Fig. 2G-I, PEG, Fig. S6).

To determine the potential sites of action for NPY-Y1R signaling in the lPBN, we first visualized neurons expressing the Y1 receptor. We found that Y1R is expressed on a subset of glutamatergic neurons in the lPBN as well as spinoparabrachial projection neurons (Fig. S7A-C). Thus, we reasoned that NPY could be acting pre- or post-synaptically on neurons in the lPBN that express the Y1 receptor. Genetic deletion of the Y1 receptor from dorsal horn projection neurons, however, did not affect hunger’s ability to suppress pain (Fig. 2J-K, Fig. S7D), suggesting that NPY likely acts post-synaptically on Y1R-expressing lPBN neurons. Together, with the pharmacological data demonstrating a role of NPY Y1R signaling in the lPBN, these results suggest that survival threats activate distinct neural populations that project to the lPBN and release NPY, which in turn acts on Y1 receptors on neurons in the PBN.

We next assessed the circuit connectivity between NPY inputs to the lPBN and Y1R expressing PBN neurons (Fig. S7E). By stimulating arcuate NPY projections to the lPBN, we determined that while a fraction of non-Y1R neurons in the lPBN receive direct synaptic inputs from NPY neurons, we did not find any evidence of synaptic connectivity between arcuate NPY neurons and lPBN Y1R neurons (Fig. S7F-H). These physiological findings are substantiated by monosynaptic rabies tracing which did not reveal direct connections between arcuate NPY and lPBN Y1R neurons (Fig. S7I-J). Taken together, these findings suggest that Y1R-expressing neurons in the lPBN could be a critical hub for the convergence of hunger, thirst, fear, and pain that dramatically influence the behavioral expression of pain. Incoming axons from multiple brain regions, are capable of influencing pain behavior by acting on Y1R-expressing neurons extrasynaptically through bulk release of NPY throughout the lPBN (Fig. 2L).

### lPBN Y1 receptor-expressing neuron activity influences sustained pain-like responses

Our findings demonstrate that NPY signaling at Y1 receptors on lPBN neurons is the likely mechanism by which competing needs suppress long term pain. To determine the functional role of lPBN Y1R neurons in pain behavior, we performed a series of neural activity manipulations. Because the Y1 receptor activates Gi-mediated signaling, we first mimicked the effect of NPY by expressing the inhibitory designer receptor exclusively activated by designer drugs (DREADD) hM4D(Gi) in lPBN Y1R neurons (Fig. 3A). We found that inhibition of Y1R neurons suppresses sustained behavioral responses to injury without influencing acute responses. Both formalin induced inflammatory pain and SNI induced mechanical hypersensitivity and cold allodynia were suppressed by DREADD inhibition of lPBN Y1R neurons (Fig 3B-C, Fig. S8A-F). Acute ongoing (Fig. 3D) and transient (Fig. 3E) nociceptive responses were not modulated by DREADD inhibition, phenocopying the effects of NPY release during competing needs. Ablating lPBN Y1R neurons also attenuated lasting pain behavior (Fig. S9A-E) without observable effects on locomotion or other aversive or threatening stimuli (Fig. S9F-O).

**Figure 3.**
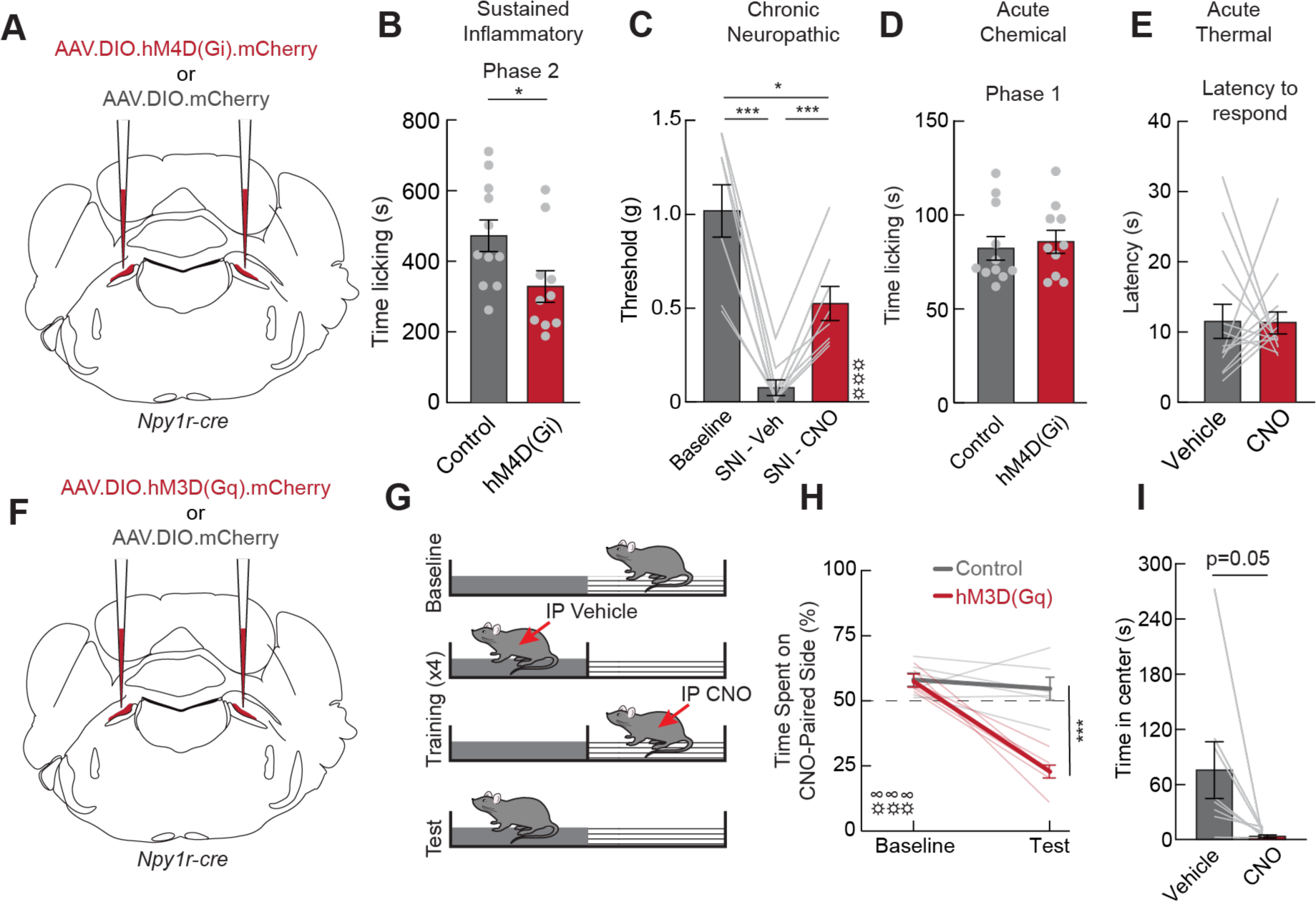
Manipulating lPBN Y1R neuron activity bidirectionally modulates pain-related behaviors. (**A**) Bilateral lPBN injections of inhibitory DREADDs in *Npy1r:Cre* mice. (**B**) Time licking paw during Phase 2 in mice injected with a control virus (grey) or with hM4D(Gi) (red). All mice were injected with CNO (1mg/kg i.p.) 15 min before formalin injection (n=10-11/group, unpaired t-test, p<0.05). (**C**) Withdrawal threshold to mechanical stimuli in hM4D(Gi)-expressing mice before SNI (baseline, grey), after SNI with a control saline injection (dark grey), and after SNI with an injection of CNO (red, n=8, repeated measures one-way ANOVA, p<0.001). (**D**) Time licking paw during Phase 1 of mice injected with a control virus (grey) or with hM4D(Gi), (red, n=10-11/group, unpaired t-test, n.s.). (**E**) Latency to respond on a 52°C hot plate in hM4D(Gi)-expressing mice treated with a saline control (grey) or CNO injection (red, n=14, paired t-test, n.s.). (**F**) Schematic depicting bilateral lPBN injections of excitatory DREADDs in *Npy1r:Cre* mice. (**G**) Schematic depicting conditioned place preference procedure. (**H**) Percent time spent in the side of the chamber paired with CNO in mice injected with a control virus (grey) or hM3D(Gq) (red) before and after conditioning (n=6-7/group, repeated measures two-way ANOVA, main effect of group p<0.001). (**I**) Time spent in the center of the open field in control and CNO treated mice (n=8, paired t-test, p=0.05). Data are expressed as mean ± SEM. Grey dots represent individual mice in between subject experiments and grey lines represent individual mice in within subjects experiments. T-test and post-hoc comparisons: *p<0.05, ***p<0.001. ANOVA main effect of group: ☼☼☼p<0.001. ANOVA interaction: ∞∞∞p<0.001.

Does activity in Y1R neurons lead to a pain-like state? To test whether Y1R neurons are sufficient to evoke behavioral and affective responses that follow injury, we expressed the excitatory DREADD receptor hM3D(Gq) in lPBN Y1R neurons (Fig. 3F). Mice strongly avoided a context that was paired with activation of lPBN Y1R neurons, suggesting that they promote negative affect (Fig. 3G-H). Activation also elicited spontaneous responses often exhibited by animals following injury such as running, jumping, and anxiety-like behavior (Fig. 3I, Fig. S8G-J). Taken together, these results suggest that lPBN Y1R neurons are an essential neural node for sustained pain and may represent a site of convergence through which pain can be regulated by endogenous circuits that prioritize survival behaviors.

### Y1R neurons are a unique subset of lPBN neurons activated during sustained pain

Our findings that modulation of lPBN neural activity selectively influenced sustained nociceptive responses led us to explore how injury changes Y1R neuron activity. We monitored neural activity dynamics in lPBN Y1R neurons during formalin-induced pain (Fig. 4A). We observed both fast fluctuations that correlated with the fine timing of licking bouts and slow, sustained increases in activity that occurred during both phases of the response to formalin administration (Fig. 4B-C). We quantified the relationship between lPBN neural activity and hindlimb licking by fitting a linear model to recorded population activity. We found that fast changes in Y1R activity were well predicted by lick bouts convolved with a fast exponential filter (Fig. 4D). We also determined that there was a residual and sustained increase in activity during both phases that persisted when animals were not actively licking (Fig. 4E). Adding a slow, behavior-independent component to the model significantly improved the fit (Fig. 4F). This suggests that Y1R population activity during the formalin assay reflects a combination of fast responses to coping behaviors on top of a slowly shifting baseline that may reflect a persistent pain state.

**Figure 4.**
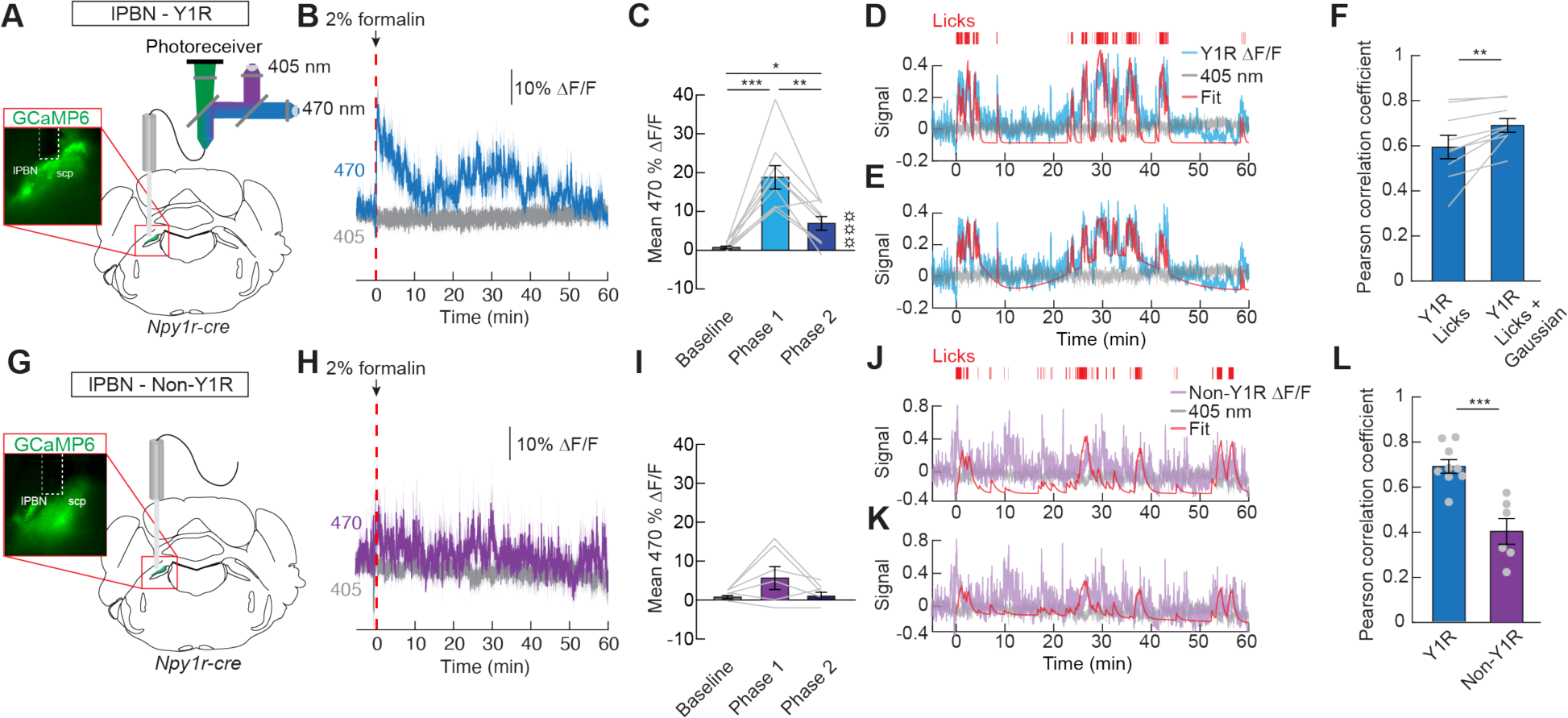
Tonic lPBN Y1R neuron activity tracks sustained pain. (**A**) Dual wavelength fiber photometry (FP) was used to measure calcium dynamics of lPBN Y1R neurons in awake, behaving mice. Inset, representative image showing GCaMP6s expression in lPBN Y1R neurons below a fiber optic tract. (**B**) Average ΔF/F of GCaMP6s signal from lPBN Y1R neurons after a formalin injection into the dorsal surface of the hindpaw (dotted red line). Signals are aligned to formalin injection. Blue, 470 nm; grey, 405 nm. Dark lines represent mean and lighter, shaded areas represent SEM. (**C**) Mean ΔF/F of lPBN Y1R neurons before formalin injection, during Phase 1, and during Phase 2 (n=9, repeated measures one-way ANOVA, p<0.001). (**D**) Representative trace showing lPBN Y1R neuron activity (blue), the control 405 nm signal (grey), licking bouts (red, above trace), and smoothed licking behavior (red trace). (**E**) Representative trace showing calcium dynamics in lPBN Y1R neurons (blue), the control 405 nm signal (grey), licking bouts (red, above trace), and smoothed licking behavior along with gaussian curves (red trace). (**F**) Pearson correlation coefficients between fit and signal for the two fitting methods (n=9, paired t-test on Fisher’s z values, p<0.01). (**G**) A cre-off construct was used to record calcium dynamics of non-Y1R expressing lPBN neurons. Inset, representative image showing GCaMP6s expression in lPBN non-Y1R neurons below a fiber optic tract. (**H**) Average ΔF/F of GCaMP6s signal from lPBN non-Y1R neurons after a formalin injection. (**I**) Mean ΔF/F of lPBN non-Y1R neurons before formalin injection, during Phase 1, and during Phase 2 (n=6, repeated measures one-way ANOVA, n.s.). (**J**) Representative trace showing lPBN non-Y1R neuron activity (purple), the control 405 nm signal (grey), licking bouts (red, above trace), and smoothed licking behavior (red trace). (**K**) Representative trace showing calcium dynamics in lPBN non-Y1R neurons (purple), the control 405 nm signal (grey), licking bouts (red, above trace), and smoothed licking behavior along with gaussian curves (red trace). (**L**) Pearson correlation coefficients between fit and signal for Y1R neurons (blue) and non-Y1R neurons (purple, n=6-9/group, unpaired t-test on Fisher’s z values, p<0.001). Data are expressed as mean ± SEM. Grey dots represent individual mice in between subject experiments and grey lines represent individual mice in within subjects experiments. T-test and post-hoc comparisons: *p<0.05, **p<0.01, ***p<0.001. ANOVA main effect of group: ☼☼☼p<0.001.

To determine whether the relationship we observed between population dynamics and behavior during sustained pain-like responses is restricted to the Y1R population of the lPBN, we monitored activity in non Y1R-expressing neurons in the same region (Fig. 4G, Fig. S10A-C). Non-Y1R neuron activity did not increase significantly after formalin administration, and the correlation between neural activity and behavior was weaker in non-Y1R neurons than Y1R neurons (Fig. 4H-L, Fig. S10D). Although other lPBN populations respond to acute nociceptive stimuli (*9, 20, 21*), the unique expression of the slow component found in Y1R population activity is further substantiated by experiments monitoring all lPBN glutamatergic neurons or other genetically defined subpopulations, none of which exhibited significant slow response components during the second (inflammatory) phase following formalin administration (Fig. S10E-P, Fig. S11).

### lPBN modulation by competing needs is consistent with a gating of nociceptive input

We have established that Y1R neurons in the lPBN are activated during sustained responses to painful stimuli and that hunger and fear suppress these responses via NPY release in the lPBN. Because our indices of pain in mice were limited to observed coping and reflexive behaviors, we used a mathematical model to infer an internal pain state based on these behaviors. We first built a reinforcement learning model based on the assumptions that animals aim to reduce pain with minimal action or “effort” and that licking is a coping behavior that reduces pain at the cost of some effort. Thus, our model system incorporates two state variables— perceived pain and exerted effort—and one available action of licking that reduces pain at the cost of effort (Fig. 5A, Fig. S12A-B). We used standard Q-learning to fit a policy (whether to lick) given the pain and effort state variables. We found that this low-dimensional state-space is sufficient to match the observed behavioral response to formalin injection in baseline conditions (Fig. 5B-D).

**Figure 5.**
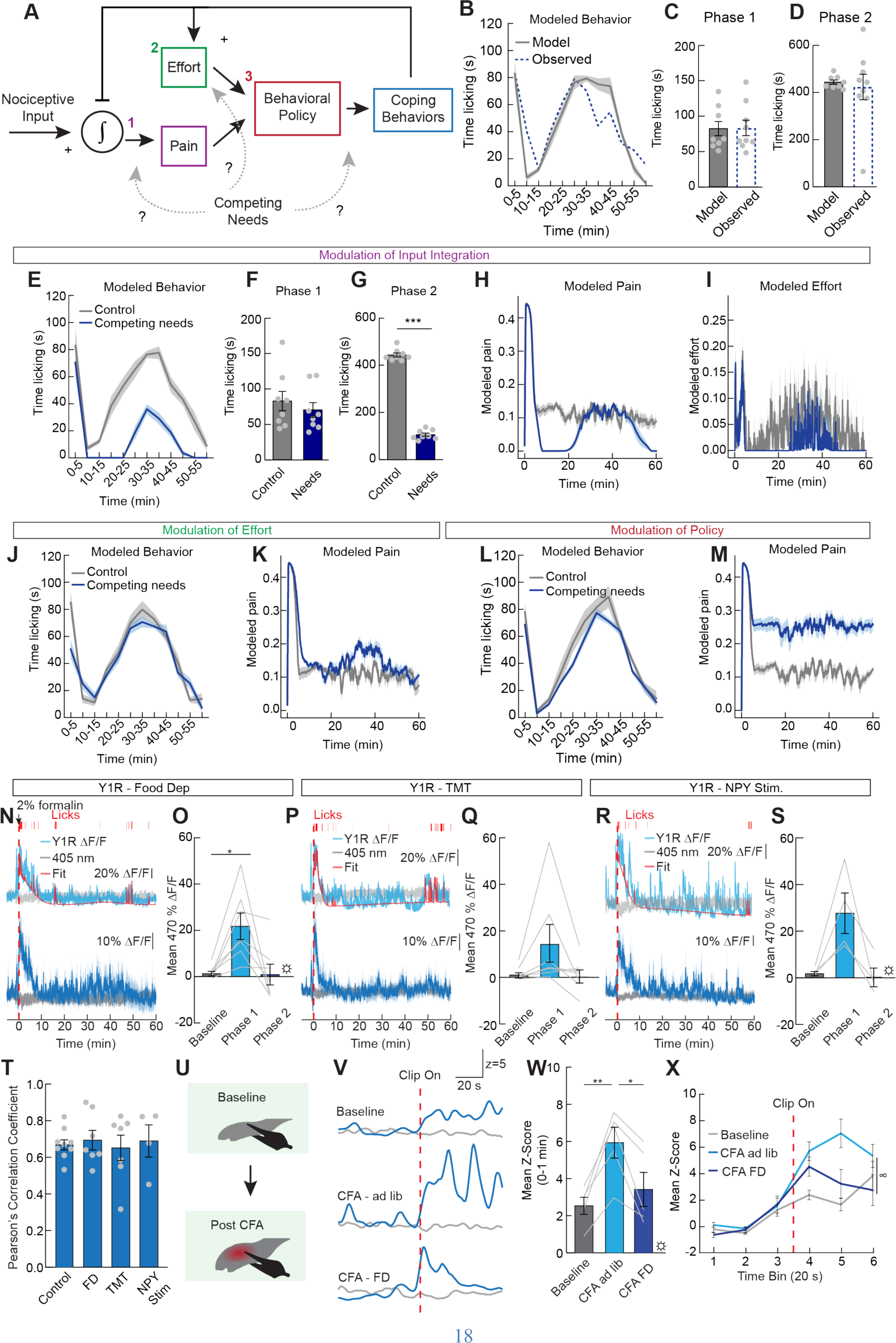
Competing drives suppress lPBN Y1R neuron activity to abrogate prolonged pain. (**A**) Schematic of the model control system for regulating behavior during pain. Nociceptive input is integrated into an internal ‘pain’ state, while past actions (licks) are integrated over recent history by an internal ‘effort’ state. The model’s learned policy maps pain and effort states to a generated behavior (lick or don’t lick) which in turn modifies the value of the two states. We study the effect of competing needs by manipulating three different parts of the model (input integration, effort, and policy, dashed arrows). (**B**) Time spent licking paw in 5 min bins of 8 trained models (grey) compared with observed mouse behavior (dotted blue). Dark lines represent mean and shaded areas represent SEM. (**C**,**D**) Time spent licking paw in phase 1 (C) and phase 2 (D). (**E**) Simulated paw licking behavior of 8 trained models, comparing control (grey) with a competing need that reduces nociceptive integration (blue). (**F,G**) Simulated time licking paw in phase 1 (F) and phase 2 (G) in the models. (**H,I**) The average dynamics of the ‘pain’ (H), and ‘effort’ (I) states of 8 trained models, comparing control with a competing need that reduces nociceptive integration (**J,K**) Simulated paw licking behavior (J) and average ‘pain’ state dynamics (K) of 8 trained models, comparing control with a competing need that increases the effort cost of licking. (**L,M**) Simulated paw licking behavior (L) and average ‘pain’ state dynamics (M) of 8 trained models, comparing control with a competing need that extends the dimension of the policy. (**N**) (top) Representative trace showing lPBN Y1R neuron activity (blue), the control 405 nm signal (grey), paw licking bouts (red, above trace), and exponential filtered licking behavior (red trace) in a food deprived mouse. (bottom) Average ΔF/F of GCaMP6s signal from lPBN Y1R neurons after a formalin injection (dotted red line) in food deprived mice. Signals are aligned to formalin injection. Blue, 470 nm; grey, 405 nm. (**O**) Mean ΔF/F of lPBN Y1R neurons before formalin injection, during Phase 1, and during Phase 2 (n=7, repeated measures one-way ANOVA, p<0.05). (**P**) (top) Representative trace showing lPBN Y1R neuron activity (blue), the control 405 nm signal (grey), paw licking bouts (red, above trace), and exponential filtered licking behavior (red trace) in mice exposed to TMT. (bottom) Average ΔF/F of GCaMP6s signal from lPBN Y1R neurons after a formalin injection in mice exposed to TMT. (**Q**) Mean ΔF/F of lPBN Y1R neurons before formalin injection, during Phase 1, and during Phase 2 (n=7, repeated measures one-way ANOVA, n.s.). (**R**) (top) Representative trace showing lPBN Y1R neuron activity (blue), the control 405 nm signal (grey), paw licking bouts (red, above trace), and exponential filtered licking behavior (red trace) in *ad libitum* fed mice with arcuate NPY neurons activated. (bottom) Average ΔF/F of GCaMP6s signal from lPBN Y1R neurons after a formalin injection with arcuate NPY neurons activated. (**S**) Mean ΔF/F of lPBN Y1R neurons before formalin injection, during Phase 1, and during Phase 2 (n=4, repeated measures one-way ANOVA, p<0.05). (**T**) Pearson correlation coefficients between fit and signal for Y1R neurons during all conditions (n=4-9/group, one-way ANOVA, n.s.). (**U**) A clip was applied to pinch the paw before and two days after an injection of CFA. (**V**) Representative trace showing lPBN ΔF/F of lPBN Y1R neurons after applying a pinch to the paw before CFA treatment (baseline), after CFA treated and *ad libitum* fed mice, and after CFA treatment and 24 h food deprived mice. (**W**) Mean z-score after the pinch is applied to the paw in the three groups shown in (V) (n=5, repeated measure one-way ANOVA, p<0.05). (**X**) Average z-score (20 s bins) during the pinch test shown in (V) (n=5, repeated measures two-way ANOVA, group x time interaction p<0.05). Data are expressed as mean ± SEM. Grey dots represent individual mice in between subject experiments and grey lines represent individual mice in within subjects experiments. T-test and post-hoc comparisons: *p<0.05. ANOVA main effect of group: ☼p<0.05. ANOVA interaction: ∞p<0.05.

We next used this model to evaluate potential ways in which a competing need signal (NPY) might impact both behavior and the underlying ‘pain state’. We performed three modulations: 1) gating the integration of ascending nociceptive input to reduce perceived pain, 2) increasing the effort “cost” of coping behavior to reduce behavioral response to pain, or 3) acting as a third “state” axis of a control policy to enable higher-dimensional mappings from state to behavior. We retrained the model using a “competing need state” condition, in which we modulated each of the three potential sites of action (Fig. 5A, Fig. S12C-D). Reducing the pain integration predicted a dramatic decrease in licking behavior during the inflammatory phase which closely mirrors the behavior observed during competing needs. The model also predicted a decrease in underlying pain state (Fig. 5E-I, Fig. S12G-I). Models of competing need states that acted on the effort cost per lick or the behavioral policy itself failed to significantly alter behavioral responses and predicted an increase, rather than a decrease, in the underlying pain state of the model during competing needs (Fig. 5J-M, Fig. S12E-F, S12J-O). This finding is consistent with NPY gating the integration of ascending nociceptive input and is not consistent with NPY reducing licking by increasing the cost of motor actions or reshaping behavioral control policy.

We noted that the model’s pain and effort states in the baseline condition resembled the slow and fast dynamics, respectively, of Y1R neural activity after formalin injection (Fig. 4B, 4D), suggesting that activity in Y1R neurons may in part reflect pain state. To determine how modeled pain compared to Y1R neuron activity during competing needs, we measured Y1R neuron responses to noxious stimuli during hunger and innate fear. Food deprivation and TMT both eliminated the Y1R neural response during the inflammatory phase of the formalin assay (Fig. 5N-Q), mirroring the observed suppression of behavioral responses. According to our model, this finding further suggests that NPY acts at the site in which nociceptive input is integrated with competing needs. The reduction in Y1R neural activity was recapitulated by chemogenetic activation of arcuate NPY (hunger) neurons that project to the PBN (Fig. 5R-S, Fig. S13). Across all conditions, nocifensive behavior and Y1R neural activity remained highly correlated (Fig. 5T). We also found that the potentiation of Y1R responses to pinching the paw during persistent inflammation was reversed by food deprivation (Fig. 5U-X), indicating that Y1R neural responses to other forms of injury that cause prolonged pain are also modulated by competing need states. In contrast, neither food deprivation nor NPY neuron stimulation affected Y1R neuron responses to acute stimuli such as noxious heat (Fig. S14), again mirroring behavioral findings and demonstrating that need states do not change the dynamics of these neurons in response to acute injury. Taken together, our modeling and neural recording experiments suggest that the slow component of Y1R neuron activity reflects an ongoing pain state that is independent of nocifensive behavior and modified by NPY signaling during competing needs.

## Discussion

Prolonged pain in the absence of an external noxious stimulus likely reflects changes in both peripheral systems and central neural networks. Our findings suggest that a variety of acute physiological and environmental needs can specifically alleviate sustained responses to injury without affecting acute nociception. We identified the parabrachial nucleus as a central hub for the convergence of needs states that allows for the gating of sustained pain through NPY release. This suggests a common mechanism is used by various survival needs to determine pain behavior. NPY release in the lPBN acts on Y1 receptor-expressing neurons to suppress pain by reducing the integration of noxious sensory input. This mechanism is an efficient and tunable system that allows urgent needs to shift brain state away from pain and towards other states that promote survival (*22*).

Deconstructing the various aspects of pain has been a challenge for centuries (*1, 4, 23*). The idea that pain is a global brain and body state rather than a transient sensory experience has become more widely accepted in recent years (*24*). Identifying, describing, and quantifying this state, especially in preclinical models, is an important step in developing treatments for pain. Assessing pain state, however, requires going beyond traditional behavioral assessments as pain states persist when animals are not actively engaged in nocifensive behaviors (*25*). For example, unbiased computational assessments of posture or facial expressions have recently been used to infer emotional states including the negative affect associated with pain (*26*). Here we find evidence for two additional indices of pain state – neurophysiological signatures of longer lasting pain states and behavioral modeling assessments based in control theory. These methods can also ultimately be used to test how pain states are affected by physiological, environmental, or therapeutic interventions. Using both neural activity recordings and modeling, we show that competing needs, through NPY release, influence the way in which lPBN Y1R neurons integrate nociceptive input to produce sensations of pain. Future work could build off these findings to test other endogenous or pharmacological interventions that curb pain state.

## Acknowledgments

We thank the Betley and Thaiss labs for comments and helpful discussions throughout the project. We thank Daniel Goldstein for help with data analysis and Grace Zhang, Jenna Golub, Hallie Kern, Callie Jaycox, Izzy Bruckman, and Mariah Wright-Moses for help with behavioral scoring.

## Funding

Klingenstein Foundation (JNB)

University of Pennsylvania School of Arts and Sciences (JNB)

National Institutes of Health grant F31DK131870 (NG)

National Science Foundation GRFP (NG)

Blavatnik Family Foundation Fellowship (NG)

## Author contributions

Conceptualization: NG, AM, AK, JNB

Data collection and analysis: NG, AM, HNA, TSN, KAK, MK, NKS, JREC, REV, EC, ELM

Visualization: NG, JNB Supervision: RK, BKT, AK, JNB

Writing – original draft and editing: NG, AM, AK, JNB

Writing – reviewing: all authors

## Data and materials availability

All data will be made available upon acceptance

## Materials and Methods

### Experimental Model and Subject Details

Mice were group housed on a 12 h light/12 h dark cycle with *ad libitum* access to food (Purina Rodent Chow, 5001) and water unless otherwise noted. Group housed adult male and female mice (at least 8 weeks old) were used for experimentation. *Npy1r-Cre* (Jackson Labs 030544, *B6.Cg-Npy1r^tm1.1(cre/GFP)Rpa^/J* (*27*), *Npy-Flp* (Jackson Labs 030211, *B6.Cg-Npy^tm1.1(flpo)Hze^/J*) (*28*), *Pdyn-IRES-Cre* (Jackson Labs 027958, B6;129S-*Pdyn^tm1.1(cre)Mjkr^*/LowlJ) (*29*), *Penk-IRES2-Cre* (Jackson Labs 025112, B6;129S-*Penk^tm2(cre)Hze^*/J), *Vglut2-IRES-Cre* (Jackson Labs 016963, *Slc17a6^tm2(cre)Lowl^*/J) (*30*), *Y1-lox/lox* (*31*), *Lbx1-Cre* (*Lbx1^tm3.1(cre)Cbm^*) (*32*), *Npy-IRES-Cre* (Jackson Labs 027851, B6.Cg-*Npy^tm1(cre)Zman^*/J) (*33*) and *C57BL/6J* mice were used for experimentation. Genotyping of the Y1-lox/lox was performed using primers and conditions provided by Jackson Labs. All other strains were genotyped by Transnetyx. All mice were habituated to handling and experimental conditions prior to experiments. For within-subject behavioral analyses, all mice received all experimental conditions. Conditions were counter balanced unless otherwise noted, and order was randomly assigned. For between-subject analyses, mice were randomly assigned to experimental condition. We performed experiments in both male and female subjects and did not observe any trends or significant sex differences. All procedures were approved by the University of Pennsylvania and University of Pittsburgh Institutional Animal Care and Use Committees.

### Recombinant Adeno-Associated Virus (rAAV) Constructs

The following rAAV vectors were used: AAV5.hSyn.DIO.hM3D(Gq).mCherry (Addgene 44361 from Bryan Roth (*34*), titer: 2.1e13 GC/mL), AAV8.hSyn.DIO.hM4D(Gi) (Addgene 44362 from Bryan Roth (*34*), titer: 2.2e13 GC/mL), AAV2.FLEX.DTR.GFP (Addgene 124364 from Eiman Azim and Thomas Jessel (*35*), titer: 2.0e13 GC/mL), AAV8.hSyn.DIO.mCherry (Addgene 50459 from Bryan Roth, titer: 3.6e13 GC/mL), AAV1.Syn.Flex.GCaMP6s.WPRE.SV40 (Addgene 100845 from Douglas Kim and the GENIE project (*36*), titer: 4.2e13 GC/mL), AAV1.hSynapsin1.axon.GCaMP6s (Addgene 111262 from Lin Tian (*37*), titer: 4.1e13 GC/mL), AAV1.syn.FLEX.splitTVA.EGFP.tTA (Addgene 100798 from Ian Wickersham (*38*), titer: 1.2e12 GC/mL), AAV1.TREtight.mTagBFP2.B19G (Addgene 100799 from Ian Wickersham (*38*), titer: 1e13 GC/mL), pSADdeltaG.mCherry (Addgene 32636 from Edward Callaway (*39*), titer: 3.8e12 GC/mL), AAV1.CBA.DO(FAS).GCaMP6s (Addgene plasmid 110135 from Bernardo Sabatini (*40*), packaged by Vigene Biosciences, titer: 4.1e13 GC/mL), AAV1.hSyn.Cre.WPRE.hGH (Addgene 105553 from James M. Wilson, 2.5e13 GC.mL) and AAV1.FLEX.tdTomato (Addgene 28306 from Edward Boyden, titer: 3.1e13 GC/mL). Syn, human Synapsin 1 promoter. FLEX, Cre-dependent flip-excision switch. WPRE, woodchuck hepatitis virus response element. GCaMP, genetically encoded calcium indicator resulting from a fusion of GFP, M13 and Calmodulin. DIO, Double-floxed inverted orientation. hM4, human M4 muscarinic receptor. DTR, diphtheria toxin receptor. Ef1a, eukaryotic translation elongation factor 1 apha. SV40, simian virus 40. TVA, tumor virus receptor A. EGFP, enhanced green fluorescent protein. HGHpA, human growth hormone polyA.

### Surgery

#### Intracerebral Viral Injections, Fiber Optic and Cannula Placement

Viral injections and implants were performed as previously described. Briefly, mice were anesthetized with isoflurane (3% induction, 1.5-3% maintenance), given ketoprofen (5 mg/kg s.c.) and bupivacaine (2 mg/kg s.c.) and placed into a stereotaxic frame (Stoelting, 51730). For viral injections, a craniotomy was performed above the injection site and virus injected at the following coordinates: lPBN: lambda −1.2 mm, midline ± 1.4 mm, skull surface −3.3 mm; subparafascicular nucleus : bregma −2.3 mm, midline ± 0.25 mm, skull surface −3.8 mm; arcuate: bregma −1.35 mm, midline ± 0.25 mm, skull surface −6.3 mm. Cre-dependent hM3D(Gq), hM4D(Gi), DTR, and mCherry were injected bilaterally (100-150 nL per side). Cre-dependent GCaMP6s, Cre-off GCamp6s, rabies helper viruses, and rabies virus were injected unilaterally (150-200 nL). Rabies helper viruses were diluted, mixed, and injected in a single solution with a final dilution of 1:200 (TVA), 1:15 (G), and (for non-specific tracing only) 1:3 (Cre). Equal volumes of Cre-off GCaMP6s and Cre-dependent tdTomato were mixed and injected in a single solution. For fiber photometry experiments, an optic fiber (400 μm core, NA 0.66, Doric) was lowered 0.2 mm above the injection site and secured with metabond (Parkell) and dental cement. For pharmacological experiments, mice were implanted with 26 gauge guide cannulae (Plastics One) above the lPBN secured with metabond and dental cement. Mice were given at least 2 weeks for recovery and transgene expression. Expression and fiber placements were verified post-mortem.

#### Spinal Cord Injections

Spinal cord viral injections were performed as previously described (*8*). Mice were anesthetized with isoflurane (3% induction, 1.5-3% maintenance) and given ketoprofen (5 mg/kg s.c.) and bupivacaine (2 mg/kg s.c.). An approximately 3 cm incision was made through the skin to expose vertebrae T12, L1, and L2. The muscle was gently removed and a laminectomy was performed to expose the spinal cord. The mouse was then placed in a stereotaxic frame to hold the spinal column in place. Cre-dependent axon GCaMP6s was injected unilaterally 0.5 mm lateral from the anterior spinal artery and 0.3 mm ventral from the dura mater. Two 500 nL injections were performed at the rostral and caudal ends of the exposed spinal cord. The skin was then sutured, and mice were given at least 3 weeks for recovery and viral expression.

### Pain Models

#### Spared Nerve Injury

Mice were anesthetized with 5% isoflurane, and anesthesia was maintained at 2% isoflurane throughout surgery. The left hindlimb was shaved with electric trimmers and sterilized with 70% ethanol and povidone-iodine (Medline). An incision (2-3 cm) was made in the skin of the upper left hindleg and the muscle was spread to expose the branches of the sciatic nerve. The peroneal and tibial branches were ligated with silk suture (6–0) and transected. The skin was closed with 9 mm wound clips and topical triple antibiotic ointment (Neosporin) was applied to the wound. Wound clips were removed approximately 10 days post-surgery. Behavioral experiments began 14 days following surgery. Sham surgery involved the same surgical procedures, including exposure of the nerve but no ligation/transection (*41*).

#### Persistent Inflammation

Complete Freund’s Adjuvant (CFA, Sigma) was diluted 1:1 in saline and subcutaneously injected (30 μl) into the plantar surface of the paw of lightly anesthetized mice (isoflurane).

### Food Deprivation

For 24 h food deprivation, mice were placed in a clean cage with alpha dry bedding and *ad libitum* access to water, but no food, 24 h prior to experimentation. *Ad libitum* fed control mice were also placed in a clean cage with alpha dry bedding but were given free access to chow. Body weight was measure before removing food and before experiments began.

### Behavioral Assays

#### DREADD-evoked food Intake

Mice were habituated for at least 1 h to a cage with a lined floor (Kimberly-Clark, 75460) and *ad libitum* access to food and water. Following habituation, food was removed from the cage and mice were injected with Clozapine-N-Oxide (CNO, 1 mg/kg i.p.) or saline and placed back in the cage with a weighed pellet of chow. Food was weighed after 1 h, accounting for crumbs.

#### Formalin Test

Mice were habituated to handling and restraint before experiments began. Mice were placed in a clear enclosure for a 15 min habituation period. Then, 20 μL of 2% formalin (Sigma) were injected subcutaneously into the dorsal surface of the hindpaw. Tests were video-recorded and time spent licking during the 1 h assay was measured by scorers blinded to experimental condition. The time spent licking was grouped into 1 or 5 min bins, and the total time spent licking during the acute (0-5 min) and inflammatory (15-60 min) phase was calculated. In experiments where unilateral lPBN recordings or infusions took place, the contralateral paw was injected.

##### Effects of food deprivation on formalin test

Food was removed 24 h prior to formalin injection. *Ad libitum* fed mice served as controls.

##### Effects of thirst on formalin test

Mice were injected with polyethylene glycol (PEG, 30% s.c.) or saline and placed back in the home cage with the water bottle removed. They were placed in an empty chamber to habituate 45 min later, and formalin was injected 60 min after PEG injection. To confirm that this dose induced thirst, a separate group of mice were injected and placed immediately into a new cage with an inverted 15 mL conical tube with a lick spout filled with water. Volume of water consumed was measured after 1 and 2 h and compared with saline injected controls.

##### Effects of predator odor on formalin test

Mice were placed in the arena with a piece of absorbent paper taped to the side for a 15 min habituation period. 30 μL of TMT (diluted 1:10 in PBS) or PBS was then pipetted onto the absorbent paper. 15 min later, formalin was injected into the paw as described above.

##### Effects of conditioned fear on formalin test

Mice were placed in a fear conditioning chamber (HABITEST modular behavioral test system, Coulbourn Instruments). After a 10 min habituation period, 1 shock (1 mA, 2 sec) was administered every 2 min over the course of 10 min (5 shocks total) (*42*). 24 h later, mice were injected with formalin as described above. Mice were placed into the environment either immediately (to measure acute phase responding) or 20 min later (to measure inflammatory phase responding). Because conditioned fear extinguishes within 10 min, pain responding was only monitored in the fear-conditioned environment for 10 min. Mice were either placed in the shocked context (square chamber, gray walls and ceiling, shock rod flooring, cleaned with 70% ethanol) or an unshocked context (rounded chamber, black and white checkered walls and ceiling, cleaned with 5% acetic acid). Behavior was video-recorded for 10 min following chamber entry. Freezing was scored automatically using ANY-Maze software (Stoelting) and manually validated. Paw licking was scored manually.

##### Effects of lPBN Y1R neuron inhibition and AgRP neuron stimulation on formalin test

CNO (1 mg/kg) was injected i.p. before the 15 min habituation period. The formalin test was then performed as described above.

#### von Frey Test

Mice were habituated to small plexiglass chambers atop mesh flooring for 2 h per day for 3 days before experimentation and for 15 min before each test. <u>The ascending method</u> (*43*): Ten von Frey filaments (ranging from 0.04 g to 10 g) were used. Starting with the smallest von Frey filament and continuing in ascending order, each filament was applied to the plantar surface of the hindpaw until the filament bent. Each filament was tested 5 times with 1 min between each trial. A positive response was defined by one or more of the following: paw withdrawal, guarding, licking, shaking, or jumping. The number of positive responses was recorded for each filament, and the percentage of responses for each filament was calculated (# of withdrawal trials/total trials). Threshold was determined as the first filament at which the mouse had a positive response to 3 or more trials. The ascending method was used for CFA experiments. <u>The up/down method</u> (*44*): Logarithmically increasing calibrated von Frey filaments (Stoelting) ranging from 0.001 g to 6.0 g were used for testing. Beginning with the 0.4 g filament, filaments were applied perpendicular to the surface of the lateral hindpaw with sufficient force to cause bending of the filament (∼30°). A positive response was defined by one or more of the following: paw withdrawal, guarding, licking, shaking, or jumping. A positive response was followed by application of a lower gram force filament, while a negative response was followed by application of a higher gram force filament to calculate the 50% withdrawal threshold for each mouse. The up/down method was used for SNI experiments.

#### Acetone-Induced Cold Sensitivity Test

Acetone drop withdrawal testing was performed in the same Plexiglas chambers following von Frey testing. A syringe attached to PE-90 tubing with a tip flared to a 3.5 mm diameter was used to apply a 10 μL droplet of acetone to the lateral surface of the hindpaw. The length of time the animal spent lifting, shaking, and attending to the paw directly following droplet application was recorded, with a 30 s cutoff. Three trials were averaged for the final result (*41*).

#### Hotplate Test

A cast iron plate with plexiglass walls was placed on a hotplate and heated to 52°C. Mice were placed on the hotplate for 1 min. All sessions were video-recorded and the latencies to lick the forepaws, lick the hindpaws, and jump were scored along with the number of jumps during the test. Scoring was performed by experimenters blinded to experimental condition.

##### Effects of predator odor on hot plate test

30 μL of TMT was pipetted onto absorbent paper taped to the plexiglass wall. The mouse was then placed on the hot plate and responses were recorded as described above. The following data points were excluded: one mouse from each group did not lick their forepaws during the assay and 2 PBS mice and one TMT mouse jumped onto the paper for several seconds and therefore their jump score was excluded.

##### Effects of food deprivation on hot plate test

Naïve mice were placed on the hot plate as described above for a habituation trial. Food was removed from half the cages and the assay was run again the next day and food was returned to all mice. The experiment was then counter balanced at least 3 days later.

##### Effects of hypothalamic AgRP/NPY neuron stimulation and Y1R neuron inhibition on hot plate test

Mice expressing DREADDs in the arcuate or PBN were placed on the hot plate as described above for a habituation trial. At least 2 days later, half the mice were injected with saline and half with CNO (i.p., 1mg/kg) followed by the hot plate test 30 min later. The experiment was then counter balanced at least 3 days later.

#### Inflammation-Induced Sensitization

An alligator clip (UQ003, Uniquers) was applied to the ventral surface of the paw as previously described to establish the baseline neural response to mechanical pain (*45*). At least two days later, CFA was injected into the plantar surface of the paw as described above. The neural response to clipping the paw was recorded 48 h later in *ad libitum* fed mice. Food was then removed from the cage. The neural response to clipping the paw was measured again, 24 h later, in food deprived mice. Paw size was measured daily using a plethysmometer (Stoelting, 57140) to ensure that any effects were not due to changes in inflammation levels.

#### Conditioned Place Avoidance

Two-sided apparatuses were used with distinct visual (black vs. white walls) and textural (flooring: plastic vs. soft textural side of Kimtech bench-top protector) cues. A neutral middle zone to shuttle between sides was maintained and the chamber was equipped with an overhead camera to track mouse position. Mice were habituated to the apparatus and a pre-conditioning preference was determined over four 30 min sessions via ANY-Maze software (Stoelting). Conditioning was performed daily for 4 days. During conditioning, mice expressing hM3D(Gq) or control virus in lPBN Y1R neurons were injected with saline (i.p.) and placed into their non-preferred side for 30 min in the morning and injected with CNO (1 mg/kg, i.p.) and placed into their preferred side for 30 min in the afternoon. To test post-conditioning preference, mice were again placed in the chamber with free access to both sides for two 30 min sessions, and their activity was tracked. The percent of time spent in the CNO-paired side of the chamber before and after conditioning was calculated and averaged to determine the shift in preference caused by activation of lPBN Y1R neurons.

#### Innate Fear Assay

Mice were placed in a chamber with absorbent paper taped to the floor on one side and their activity was tracked by an overhead camera. After a 15 min baseline period, 30 μL of TMT was pipetted onto the paper and mice were left in the chamber for another 15 min. Using AnyMaze, we quantified the number of entries into and time spent in the “odor zone,” which was defined as the area around the absorbent paper with a circumference of the mouse’s body length (excluding the tail).

#### Conditioned Taste Aversion

Mice were habituated to cages with lined flooring for at least 1 h and food was removed from the home cage for 24 h as described above. On day 1, mice were given 0.5 g of strawberry or orange jello. 5 min after their first bite, mice were injected with either saline (control) or LiCl (to induce aversion, 125 mg/kg, i.p.) and allowed to finish the jello. Mice were then returned to their home cages with *ad libitum* access to chow and water. The next day, food was again removed from the cage. On the third day, mice were presented with 1.5 g of the same flavor of jello they had on day 1, and the weight of the remaining jello was recorded at 30 min and 60 min.

#### Open Field Assay

Mice were habituated to the open field area (400mm^2^) for 15 min. Mice were then tracked by ANY-Maze software (60 min for ablation experiments, 30 min for all other experiments). The center was defined as the middle third of the arena.

### Drugs and Pharmacology

Formalin (2%, Sigma), PEG (30%, average M.W. 1500, Acros Organics), CFA (50%, Sigma), LiCl (12.5 mg/mL, Sigma), CNO (0.1 mg/mL, Tocris) and Diphtheria toxin (DT, 0.5mg/ml, List Labs) were diluted in normal saline. Formalin and CFA were injected subcutaneously in the hindpaw at a volume of 20 μL (formalin) or 30 μL (CFA). PEG was injected subcutaneously at a volume of 20 μL/g body weight. LiCl and CNO were injected intraperitoneally at a volume of 10 μL/g body weight. DT was injected intramuscularly at a volume of 5 μL/g body weight. Mice received 3 DT injections under light isoflurane anesthesia with 48 h between injections.

#### Effects of lPBN NPY Y1 receptor antagonist on the inhibition of inflammatory pain by innate fear

For all experiments, mice were habituated to handling and infusion procedures. BIBO 3304 (Tocris) was dissolved in DMSO and frozen in aliquots. Aliquots were thawed and diluted 1:1 in artificial cerebrospinal fluid (aCSF) before each experiment. 3 μg of BIBO 3304 or vehicle (1:1 solution of DMSO and aCSF) was microinjected (100 nL) with a Hamilton syringe attached to an internal cannula (Plastics One) and microliter syringe pump (PHD Ultra, Harvard Apparatus) into the lPBN of mice immediately before the formalin test (see above).

#### Effects of lPBN NPY on persistent inflammatory pain

Neuropeptide Y (NPY, Tocris) was dissolved in saline and frozen in aliquots. Aliquots were thawed and diluted 1:1 in artificial cerebrospinal fluid (aCSF) before each experiment. 0.1 μg of NPY (100 nL) or vehicle (aCSF, 100nL) was microinjected with a Hamilton syringe attached to an internal cannula as described above. Mice were then placed on the caged floor and allowed to habituate for 15 min before the von Frey test began.

### Calcium Imaging

#### Fiber Photometry (FP)

Dual-wavelength fiber photometry was performed as previously described (*46, 47*). Two excitation wavelengths (470 nm and 405 nm) were delivered through fiber-coupled LEDs (Thorlabs, M470F3 and M405F1) modulated at 211 and 566 Hz by a real-time amplifier (Tucker-Davis Technologies, RZ5P or RZ10x). Excitation lights were filtered and combined by a fluorescence minicube (Doric). The light was delivered through a 400 µm core, 0.57 NA low autofluorescence optical fiber (Doric, MFP_400/430/1100-0.57_1m_FC-MF2.5_LAF) to the implanted fiber (Doric, MFC_400/430-0.66_4.0mm_MF2.5_FLT). The fibers were secured by a clamp (Thorlabs, ADAF2). LED power was set between 20 and 40 µW emitted from the fiber tip to minimize bleaching. GCaMP fluorescence was collected from the same optical fiber and focused onto a femtowatt receiver (Newport, Model 2151, gain set to DC LOW). Fluorescence was sampled at 1017 Hz and demodulated by the processor. LEDs were externally controlled by Synapse (Tucker-Davis Technology) and synchronized cameras (Ailipu Technology) were used to video-record mice during experiments. Mice were habituated to tethering before experiments began. A 5 min baseline before any stimulus was given was recorded and used to normalize subsequent changes in fluorescence.

#### Fiber Photometry Analysis

Data were exported to MATLAB (MathWorks) from Synapse using scripts provided by Tucker-Davis Technology. Custom MATLAB scripts were used to independently normalize the demodulated 470 nm and 405 nm signals. ΔF/F was calculated (F-F_baseline_)/F_baseline_, with F_baseline_ being the median of the 300 s before the stimulus. Z scores were used in cases where multiple days separated within subject recordings in order to correct for changes in signal strength over time. Data were down-sampled to 1 Hz. Subsequent plotting and analyses were performed in MATLAB and Prism 10 (GraphPad). Mean ΔF/F was calculated by integrating ΔF/F over a period of time after the stimulus and then dividing by the integration time.

#### Fitting FP signal to behavior

In order to fit the photometry signal to licking behavior, we denote the ΔF/F photometry signal as *φ*(*t*). First, we subtract the value such that one percent of the signal *φ*(*t*) is negative and fit the behavior of each individual mouse to its photometry recording by minimizing the sum of squares:

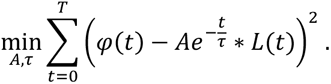

*L* is the licking behavior, a binary sequence, and it is convolved with an exponential kernel. We find the parameters by looping over time constant *τ*, and utilize least squares to find *A*. To constrain *τ*, we choose the time constant which gives the highest average Pearson correlation on all other mice in the same experimental condition. We report the ensuing Pearson correlation coefficients, indicating the extent to which the behavior co-varies with the bulk calcium signal. We model the slow component, denoted by *s*(*t*), in the photometry signal by two Gaussian curves:

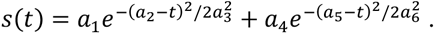

We fit the six parameters to the lower envelope of the calcium signal using MATLAB functions *envelope* and *fmincon*. Specifically, we subtract the lower envelope using argument ‘peak’ and a smoothing of 30 seconds. We do this to model the slow baseline signal on top of which the fast behavioral fluctuations ride. We then fit the behavioral sequence *L*(*t*) to the difference between the photometry signal and the slow component, using the earlier described method:

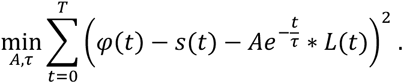

### Model of Pain Behavior

To model the control of pain-evoked licking behavior under competing needs, we assume that animals aim to reduce experienced pain while also minimizing effort expended (*48*). Animals show multiple behaviors in response to acute and chronic pain, including active coping methods like wound licking (an instinctive behavior that cleans and disinfects wounds (*49*)) and more passive strategies such as paw guarding to reduce allodynia; for the purpose of comparison to experimental data, this model focuses only on wound licking behavior. Excess wound licking leads to risk of lesion formation and infection (*50*), and the act of wound licking costs energy and saliva and comes with an “opportunity cost”, in that it prevents the animal from pursuing other activities required for survival. “Effort” in this model may therefore be taken as an abstract variable that encapsulates the overall negative consequences of licking more than is necessary to reduce pain.

We therefore posit a minimal reinforcement learning model of pain-related behavior consisting of two state variables and one action variable, where the action is either licking (*L* = 1) or not licking (*L* = 0), and the state variables are “pain” (P) and “effort” (E). Mathematically, the model aims to maximize the sum of discounted future reward, which in a model aimed at minimizing pain and effort is inversely proportional to the sum of these two quantities squared, inspired by Keramati & Gutkin (*51*)

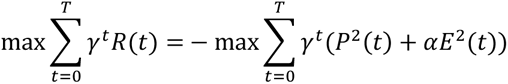

where *R_t_* is the reward at time t and T is the end of the simulation. The parameter *γ* is smaller than 1 and determines temporal discounting. The parameter *⍺* may set the relative cost of pain vs. effort but is here chosen to be one. Given this definition, a behavioral policy is defined as a mapping from the two-dimensional state of the system (*P*(*t*), *E*(*t*)) to the action *L*(*t*) (whether to lick).

Before finding this behavioral policy, we must first define how the two state variables—pain and effort—are modified by the licking action L and by ascending nociceptive input in the assay. We first consider the effort state variable, denoted E. Effort is exerted with each lick, and gradually recovers after a lick bout is over; it therefore takes the form of an exponential filter of the licking:

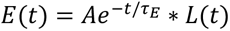

where the star ∗ denotes the convolution operation, *τ_E_* is the time constant of exponential smoothing and *A* is the scaling factor. As an abstract state variable, the units of E are somewhat arbitrary. Therefore, to select the values of *τ_E_* and *A*, we referenced the range of values found for exponential and scaling terms when fitting a linear filter of licking behavior and slow input to the photometry data. This gave us values of *τ_E_* = 4 *sec* and *A* = 0.07. We next considered the pain state variable, denoted P. In designing the form of P, we noted that a linear model could not reproduce the finding that competing need states inhibit inflammatory but not acute pain, unless we either 1) assume these two forms of pain come from different sources, one of which is independent of regulation by NPY, or 2) introduce a nonlinearity in the relationship between ascending pain and the pain state. To test whether the latter model of a single pain source would be sufficient to account for observed behavior, we here introduced a nonlinearity into P, designing it to be a saturating integral of the ascending nociceptive input from the formalin injection:

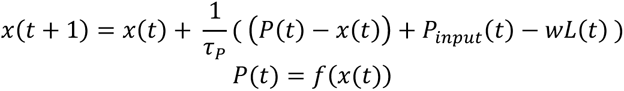

where the subscript denotes time, *τ_P_* is an integration time constant, *w* is the degree to which licking affects the pain state, and *P*_input_ is ascending nociceptive input. Like E, P is an abstract quantity with arbitrary units, thus we set our integration time constant and licking weight to ensure P and E were of comparable magnitude: this gave us *w* = 0.035 and *τ_P_* = 5 *sec*. For the case of formalin injection, *P*_input_ was modeled as the sum of two Gaussians, such that P resembles the slow signal in the photometry (see *Fiber photometry analysis* and supplementary Figure 12. *f*(.) is a nonlinearity that keeps the value of P between a minimal and maximal level (see supplementary Figure 12), {*f*(*x*) = 0 if *x* < 0, *f*(*x*) = *x* if *x* < *K* and *f*(*x*) = *K* + *tanh*(10*x*)/10 if *x* > *K*}. Here, the value K is chosen to match model output to mice *ad libitum* licking behavior, K=0.35. Thus, x is an intervening variable shaped by 1) the effect of licking, 2) the ascending pain input, and 3) the difference between x and P, which is positive when x<0, 0 in the linear portion of *f*, and negative when x>K. The difference term therefore has the effect of a restoring force that reduces excursions outside the linear regime of *f*.

Having established E and P, we can next learn a behavioral policy that maximizes reward (that is, minimizes 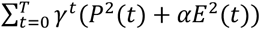.) We parametrized the model’s behavioral policy by a feedforward network: *L*(*t*) = *g*_*θ*_(*P*(*t*), *E*(*t*)), and used Q-learning (*52*) to find the network parameters *θ*. The feedforward network is all-to-all connected, with six hidden layers of 128 rectified linear units. To avoid having to model the duration of lick bouts, when the policy generates a lick, the duration is sampled from the empirical distribution of lick bout durations found in the data.

We study the effect of competing needs in the model by modulating the three parts of the model as shown in Figure 5A, denoting the modulation by a constant NPY>0. During training, 25% of trials were randomly selected to include the modulation and in other trials we set NPY=0. Each of the three parts is manipulated in two ways.

#### Modulation I: modeling a change in the integration of pain input

We tested two alternative ways of manipulating integration of nociceptive input:

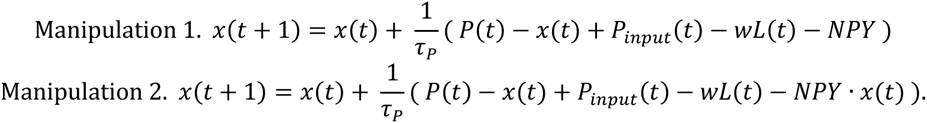

We find that this model can reproduce the experimentally observed reduction in licking when we set NPY=0.001 and NPY=0.0075 respectively.

#### Modulation II: modeling an increase in the effort cost of licking

We tested two alternative ways of manipulating the effort cost of licking, either by doubling the time constant *τ*_E_ or doubling the amplitude *A* in the equations for the dynamics of E:

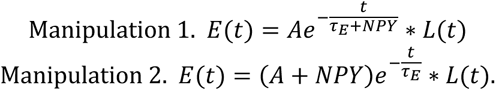

#### Modulation III: changing the behavioral policy

We tested two alternative ways of incorporating the third axis to the behavioral policy, introducing NPY as a scaling of either the pain variable or the effort variable in the reward function:

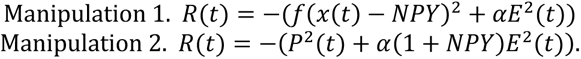

Here, NPY=0.1 and NPY=0.2 respectively. In this revised policy, NPY is given as an additional input to the feedforward network: *L*(*t*) = *g*_*θ*_(*P*(*t*), *E*(*t*), *NPY*), and we assume that its value remains constant during the time of the experiment.

Only the modulation to pain state integration leads to behavioral and state timeseries consistent with the data. We show the first manipulation of each modulation in Figure 5, while the second manipulation is shown in supplementary Figure 12. The code will be made publicly available upon acceptance.

### Histology

#### Verification of viral expression and fiber placement

Mice were transcardially perfused with 0.1 M phosphate buffered saline (PBS) followed by 4% paraformaldehyde (PFA). Brains were removed and post-fixed for 4 h in PFA and then washed overnight in PBS. Coronal brain sections were cut (150-200 μnm sections) on a vibratome, washed twice with PBS, and mounted and coverslipped with Fluorogel. Epifluorescence images were taken on a Leica stereoscope to verify fiber and cannula placements and viral expression. In rare cases (1 Y1R, 2 Vglut2, 1 non-Y1R) animals died before fiber locations could be determined.

#### Immunohistochemistry

Brain sections were incubated overnight at 4°C with primary antibodies diluted in PBS, 1% BSA and 0.1% Triton X-100. Antibodies used: goat anti-AgRP (1:2,500, Neuromics, GT15023), rabbit anti-cFos (1:1500, Cell Signaling Technology, 2250) guinea pig anti-RFP (1:10,000) (*53*), and rabbit anti-GFP (1:5,000, Invitrogen, A-11122).

Sections were washed 3 times and incubated with species appropriate and minimally cross-reactive fluorophore-conjugated secondary antibodies (1:500, Jackson ImmunoResearch) for 2 h at room temperature. Sections were washed twice with PBS and mounted and coverslipped with Fluorogel. Confocal images were taken on a Leica DM6 Upright Microscope using a 20X or 63X objective.

#### Fluorescence in situ hybridization (FISH)

Brains were post-fixed overnight and then transferred to a 30% sucrose solution for 48 h. Brains were then frozen in OCT compound (Fisher Scientific) and stored at −80°C. Brains were sectioned at 12-15 µm in a cryostat onto Superfrost Plus slides (Fisher Scientific) and stored at −80°C until use. FISH was performed according to manufacturer instructions (RNAscope fixed-frozen tissue, ACD) using the following probes: Mm-Npy1r (427021) and Mm-Slc17a6 (319171).

##### FISH Imaging

Sections were imaged on a slide scanning light microscope (Keyence). Exposure times were kept consistent across all sections in an experiment. Tile scans were collected of the entire lPBN using a 40X objective and stitched using BZ X800 Analyzer Software (Keyence).

##### FISH Analysis

FISH analyses were performed in ImageJ (NIH). ROIs were drawn manually by circling all DAPI-positive areas in the lPBN. Then, in each channel, the intensity of 5 representative “background” regions without any fluorescent labeling and the intensity of 20 single “dots” representing an mRNA transcript were measured. The approximate dot number per cell was calculated by dividing the total intensity of a DAPI-labeled ROI minus the background by the average dot intensity. A cell was considered positive if it had an estimated dot number greater than or equal to 1.

### Slice Electrophysiology

#### Slice preparation

Animals were deeply anesthetized with isoflurane and rapidly decapitated. Following brain dissection, the brain was submerged in oxygenated ice-cold N-methyl-D-glucamine (NMDG) recovery solution (in mM: 2.5 KCl, 20 HEPES, 1.2 NaH_2_PO_4_, 25 Glucose, 93 NMDG, 30 NaHCO_3_, 5.0 Sodium ascorbate, 3.0 sodium pyruvate, 10 MgCl_2_, and 0.5 CaCl-2H_2_0; 300-305 mOsm). The brain was blocked and coronal slices were obtained using a Campden Instruments 7000 smz-2 vibratome in ice-cold, oxygenated NMDG solution. After sectioning, slices were transferred to a recovery holding chamber containing 32° C NMDG solution. Slices recovered for 10 min before being transferred to a holding chamber containing room temperature artificial cerebrospinal fluid (ACSF, in mM: 119 NaCl, 2.5 KCl, 1.3 MgCl_2_-6H_2_O, 2.5 CaCl_2_-2H_2_O, 1.00= NaH_2_PO_4_-H_2_O, 26.2 NaHCO_3_, and 11 glucose; 287-295 mOsm). Slices were allowed to recover for an additional 1 h prior to recording.

#### Electrophysiology

Electrophysiology experiments were performed on a Scientifica Slicescope Pro system. Recordings were acquired using a Multiclamp 700B amplifier and a Digidata 1550B digitizer. Data was filtered at 2 kHz and digitized at 10 kHz. Recordings took place in the lateral parabrachial nucleus dorsal to the superior cerebellar peduncle and ventral to the ventral spinocerebellar tract. Y1R-expressing neurons were identified by expression of the GFP reporter and patched with 3-6 MΩ pipettes prepared with a Sutter Instrument P-97 pipette puller and containing a CsMeSO3 based internal solution (in mM: 120 CsMeSO_3_, 15 CsCl, 8 NaCl, 10 HEPES, 0.2 EGTA, 5 TEA-Cl, 4 Mg_2_-ATP, 0.3 Na2-GTP, 0.1 Spermine tetrahydrochloride, 5 QX-314 bromide, 5 Phosphocreatine disodium salt; pH 7.4, 290 mOsm). Membrane potential was clamped at −60 mV and cells were allowed to equilibrate for 5 min following establishment of the whole-cell configuration. Channelrhodopsin-mediated optically-evoked post synaptic potentials were generated using 0.3 ms 490 nm light pulses from a CoolLED pE-4000 illumination system.

### Quantification and Statistical Analysis

Data were expressed as means ± SEMs in figures and text. Paired or unpaired two-tailed t-tests and Pearson regressions were performed as appropriate. One-way, two-way, and three-way ANOVA were used to make comparisons across more than two groups using Prism 10 (GraphPad). To compare Pearson’s correlations between groups, a Fisher’s z transformation was performed on the r values before running t-tests. Equal variance was not assumed, and Welch’s corrections or Gessier-Greenhouse corrections were used where appropriate. Test, statistics, significance levels, and sample sizes for each experiment are listed in Supplementary Tables 1 and 2. ns p>0.05, t-tests and post-hoc comparisons: *p<0.05, **p<0.01, ***p<0.001; interaction: ∞p<0.05, ∞∞p<0.01, ∞∞∞p<0.001; main effect (group, condition or drug): ☼<0.05, ☼☼p<0.01, ☼☼☼p<0.001.

## Supplementary Text

While hunger, thirst, and conditioned fear only suppress the second phase of formalin pain, innate fear (TMT) attenuates both the second and first phases (Fig. S3B). We find, however, that only the second phase of formalin is rescued with microinfusion of the Y1 antagonist into the PBN (Fig. 2B-D). These results suggest that the suppression of the first phase by TMT is likely mediated by a distinct mechanism, potentially via descending modulation that has been implicated in stress induced analgesia during acute pain stimuli (*54–56*). Indeed, in our fiber photometry experiments we noted reduced Y1R neural activity during Phase 1 which would be consistent with decreased spinal activation (Fig. 5P-Q). We also note that it is unlikely that freezing behavior or immobility is simply preventing animals from displaying pain behaviors for three reasons. First, responses in the hot plate assay are not affected by TMT exposure (Fig. S3E-H). Second, conditioned fear did not alter paw licking behavior during the acute phase (Fig. S3D). Finally, despite increasing paw licking, the Y1R antagonist did not change time mobile during the formalin assay (Fig. S5A).

**Figure S1.**
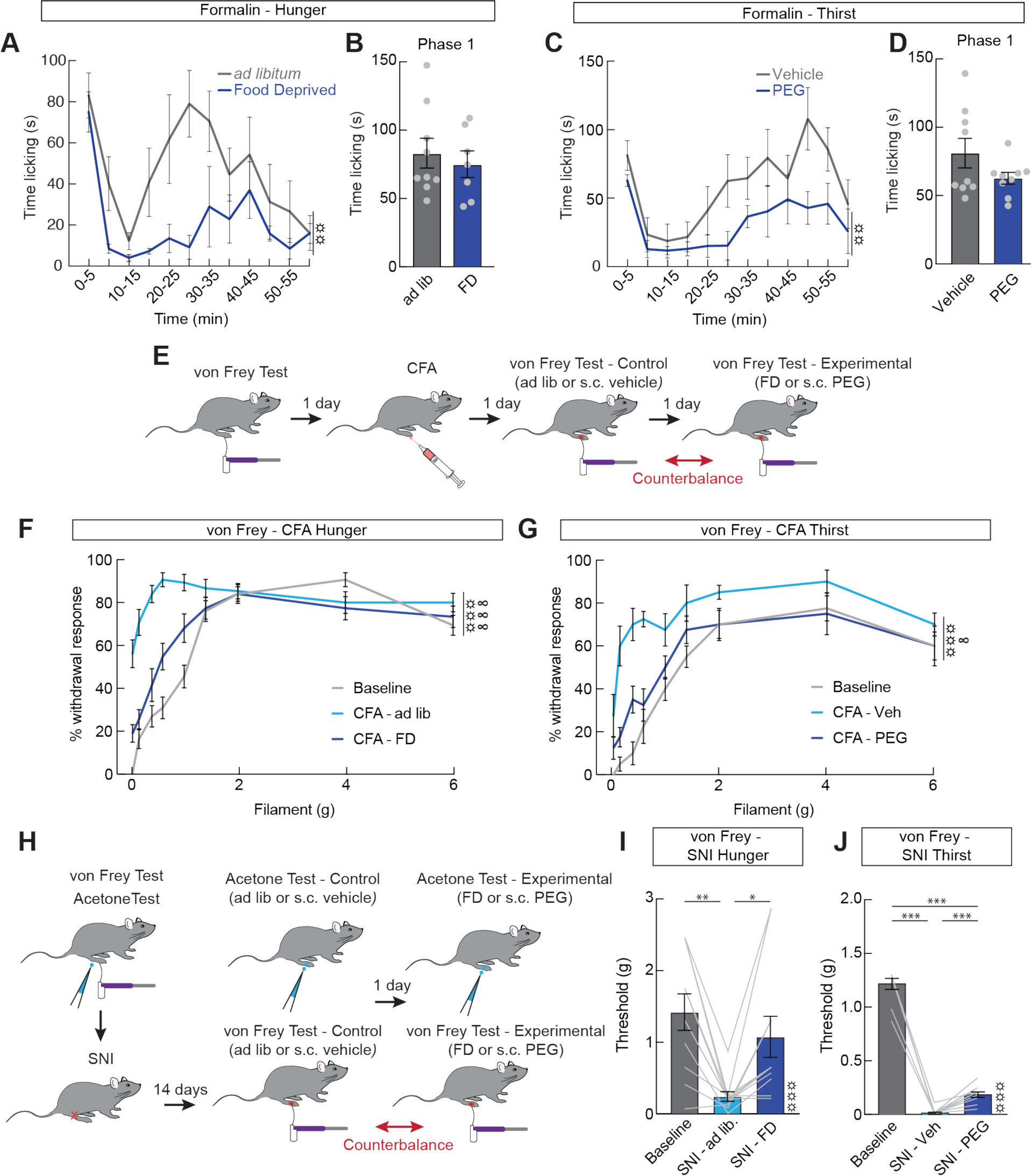
Acute nociception is not affected by physiological competing needs. (**A**) Time spent licking paw in 5 min bins after an injection of formalin in *ad libitum* fed (grey) or 24 h food deprived (blue) mice (n=7-9/group, repeated measures two-way ANOVA, main effect of group p<0.01). (**B**) Time spent licking paw during Phase 1 (n=7-9/group, unpaired t-test, n.s.). (**C**) Time spent licking paw in 5 min bins after a control injection of saline (grey) or PEG (blue, n=9/group, repeated measures two-way ANOVA, main effect of group p<0.01). (**D**) Time spent licking during Phase 1 (n=9/group, unpaired t-test, n.s.). (**E**) Experimental timeline to test the effect of food deprivation and thirst on mechanical hypersensitivity induced by persistent inflammation. Food deprived/*ad libitum* and vehicle/PEG trials were counter balanced. (**F**) Percent of trials where a paw withdrawal was observed after mechanical stimulation with the von Frey filament before CFA (baseline, grey), after CFA in *ad libitum* fed mice (light blue), and after CFA in food deprived mice (dark blue). Each von Frey filament was applied 5 times (n=15, repeated measures two-way ANOVA, main effect of group p<0.001). (**G**) Percent of trials where a paw withdrawal was observed after stimulation with the von Frey filament before CFA (baseline, grey), after CFA in control saline treated mice (light blue), and after CFA in PEG treated mice (dark blue, n=8, repeated measures two-way ANOVA, main effect of group p<0.001). (**H**) Experimental timeline to test the effect of food deprivation and thirst on mechanical and cold hypersensitivity induced by neuropathic injury. Food deprived/*ad libitum* and vehicle/PEG trials were counter balanced. (**I**) Withdrawal threshold before SNI (grey), after SNI in *ad libitum* fed mice (light blue), and after SNI in food deprived mice (dark blue, n=11, repeated measures one-way ANOVA, p<0.001). (**J**) Withdrawal threshold before SNI (grey), after SNI in control saline treated mice (light blue), and after SNI in PEG treated mice (dark blue, n=10, repeated measures one-way ANOVA, p<0.001). Data are expressed as mean ± SEM. Grey dots represent individual mice in between subject experiments and grey lines represent individual mice in within subjects experiments. T-test and post-hoc comparisons: *p<0.05, **p<0.01, ***p<0.001. ANOVA main effect of group: ☼☼p<0.01, ☼☼☼p<0.001. ANOVA interaction: ∞p<0.05, ∞∞∞p<0.001.

**Figure S2.**
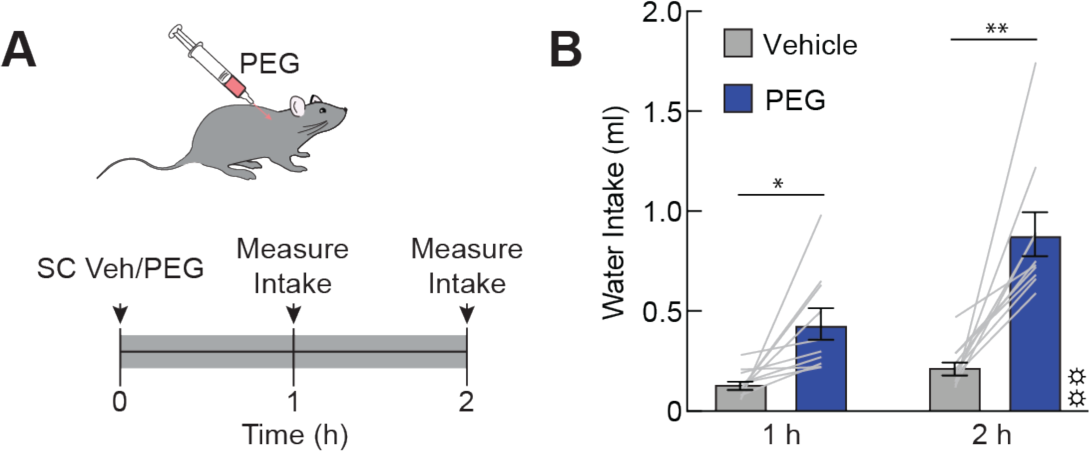
Validation of PEG-induced drinking behavior. (**A**) Water intake in control saline or PEG (30%, 20ul/g body weight s.c.) treated mice was assessed to validate thirst experiments. (**B**) Water intake 1 and 2 h after control saline (grey) or PEG (blue) treatment (n=10, repeated measures two-way ANOVA, main effect of group p<0.01). Data are expressed as mean ± SEM. Grey lines represent individual mice. Post-hoc comparisons: *p<0.05, **p<0.01. ANOVA main effect of group: ☼☼p<0.01.

**Figure S3.**
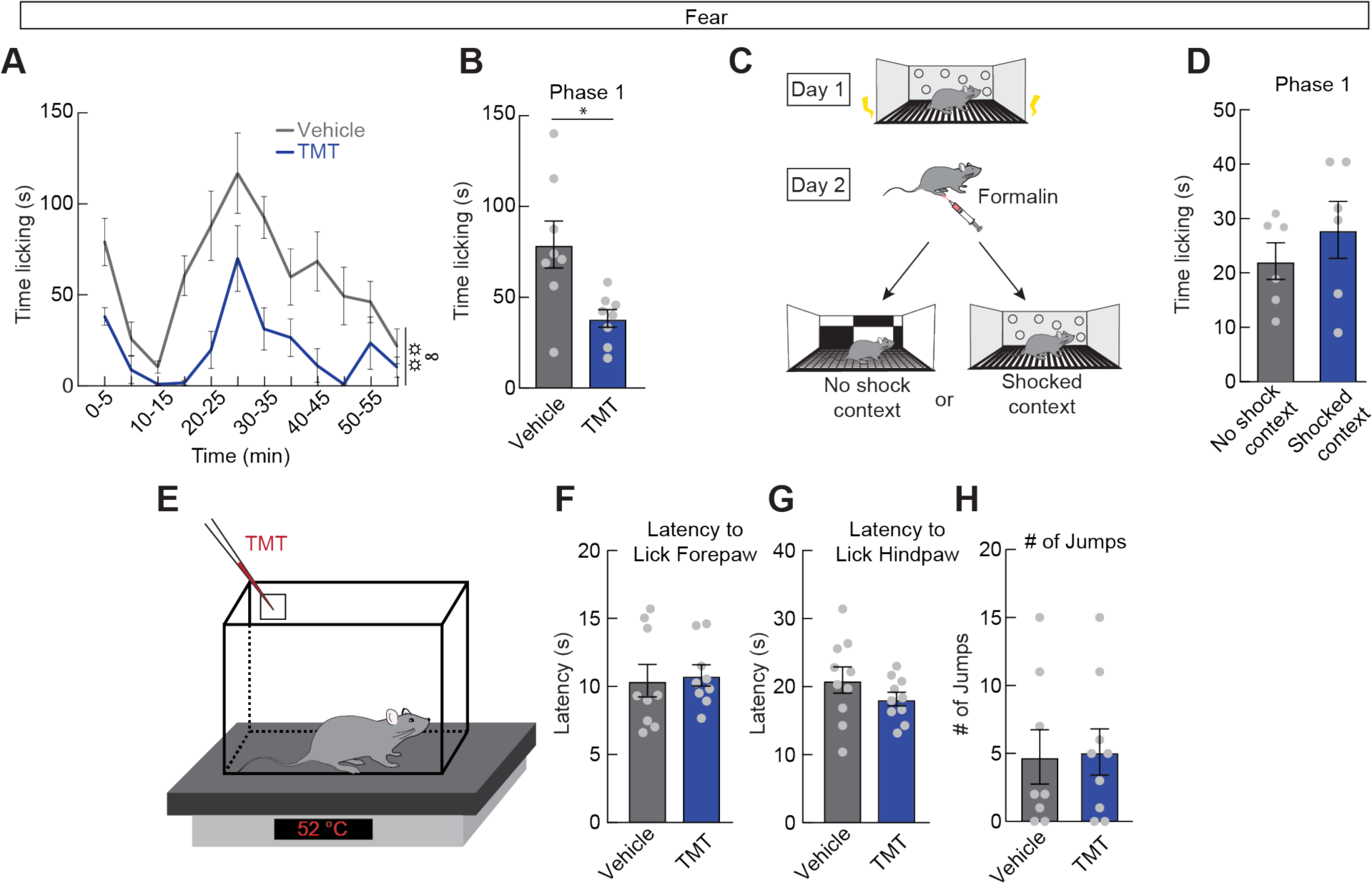
Acute nociception is not systematically affected by fear. (**A**) Time spent licking paw in 5 min bins with control PBS in the chamber (grey) or TMT in the chamber (blue, n=8/group, repeated measures two-way ANOVA, main effect of group p<0.01). (**B**) Time spent licking paw during Phase 1 (n=8/group, unpaired t-test, p<0.05). (**C**) Experimental timeline to test the effect of conditioned fear on acute formalin pain. Conditioning occurred on day 1. On day 2, mice were injected with formalin and then immediately placed in either the shocked context or a novel, unshocked context. Licking behavior was scored for 5 min. (**D**) Time spent licking paw in either the unshocked context (grey) or shocked context (blue, n=6/group, unpaired t-test, n.s.). (**E**) TMT was applied to a chamber placed on 52°C hot plate. Mice were then placed in the chamber and their behavior was video recorded for 1 min. (**F-H**) Latency to lick the forepaw (F) and hindpaw (G) and the number of jumps (H) on the hot plate after application of TMT or control PBS (n=8-10/group, unpaired t-tests, all n.s.). Data are expressed as mean ± SEM. Grey dots represent individual mice. T-test and post-hoc comparisons: *p<0.05. ANOVA main effect of group: ☼☼p<0.01. ANOVA interaction: ∞p<0.05.

**Figure S4.**
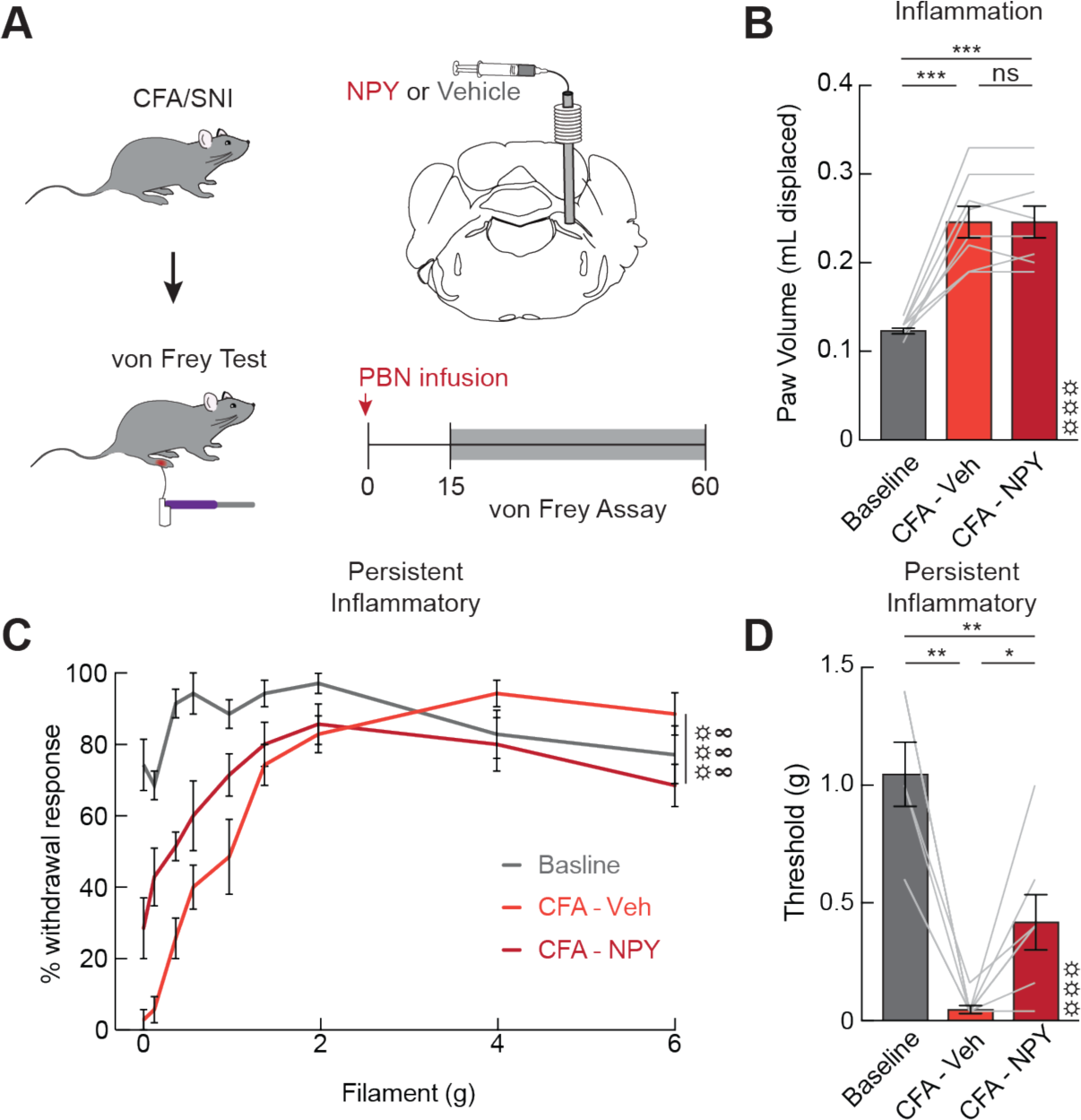
lPBN NPY signaling suppresses inflammation-induced mechanical allodynia. (**A**) Experimental timeline to test the effects of lPBN NPY signaling on inflammatory pain. Vehicle and NPY trials were counter balanced. (**B**) Paw volume before CFA injection (baseline, light grey), after CFA with a vehicle infusion (orange), and after CFA with an infusion of NPY (red, n=8, repeated measures one-way ANOVA, p<0.001). (**C**) Percent of trials where a paw withdrawal was observed after stimulation with the von Frey filament before CFA (baseline, grey), after CFA in vehicle treated mice (orange), and after CFA in NPY treated mice (red). Each filament was applied 5 times (n=7, repeated measures two-way ANOVA, main effect of group p<0.001). (**D**) Withdrawal threshold to mechanical stimuli before CFA (baseline, grey), after CFA with a vehicle infusion (orange), and after CFA with an infusion of NPY (red, n=7, repeated measures one-way ANOVA, p<0.001). Data are expressed as mean ± SEM. Grey lines represent individual mice. T-test and post-hoc comparisons: *p<0.05, **p<0.01, ***p<0.001. ANOVA main effect of group: ☼☼☼p<0.001. ANOVA interaction: ∞∞∞p<0.001.

**Figure S5.**
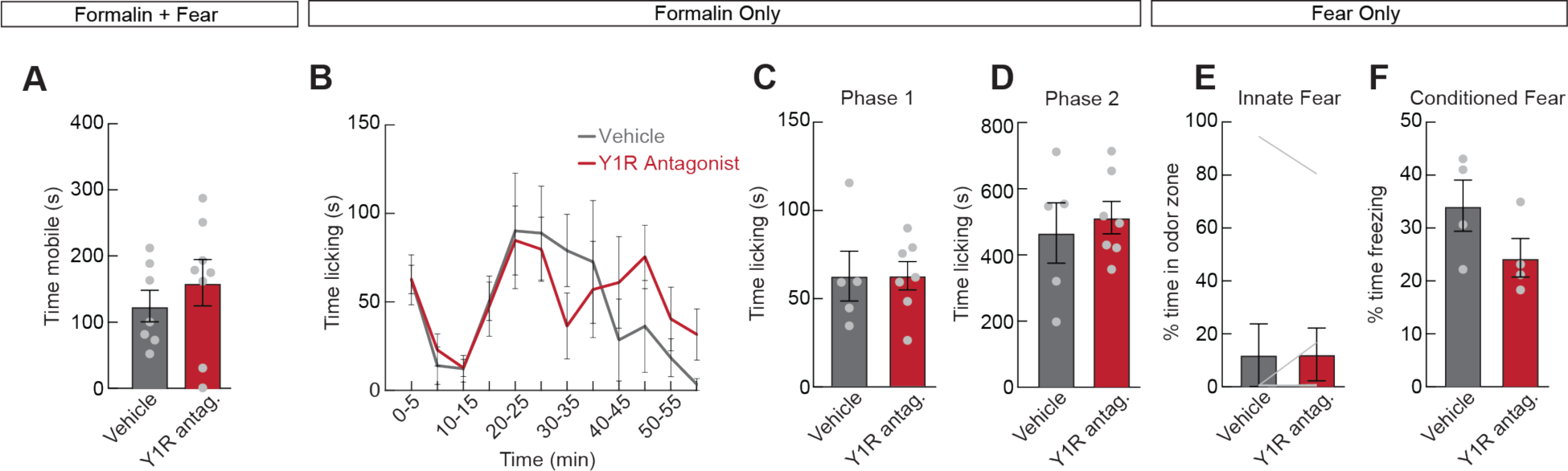
Control experiments demonstrating that lPBN Y1R signaling does not change fear responses or baseline nociceptive thresholds. (**A**) Time spent mobile during the formalin assay in the presence of TMT after a control vehicle infusion (grey) or Y1R antagonist (red, n=7-8/group, unpaired t-test, n.s.). (**B**) Time spent licking paw in 5 min bins after an injection of formalin and an infusion of vehicle (grey) or Y1R antagonist (red, n=5-7/group, repeated measures two-way ANOVA, n.s.). Blocking the Y1 receptor does not potentiate pain responses, suggesting that NPY only functions to modulate pain during competing needs. (**C**) Time spent licking paw during phase 1 (n=5-7/group, unpaired t-test, n.s.) (**D**) Time spent licking paw during phase 2 (n=5-7/group, unpaired t-test, n.s.) (**E**) Time spent in the odor zone after a control infusion of vehicle (grey) or Y1R antagonist (red) in TMT exposed mice (n=8, paired t-test, n.s.). (**F**) Percent time spent freezing after a control infusion of vehicle (grey) or Y1R antagonist (red) when mice were placed in a chamber associated with shock (n=4/group, unpaired t-test, n.s.). Data are expressed as mean ± SEM. Grey dots represent individual mice in between subject experiments and grey lines represent individual mice in within subjects experiments.

**Figure S6.**
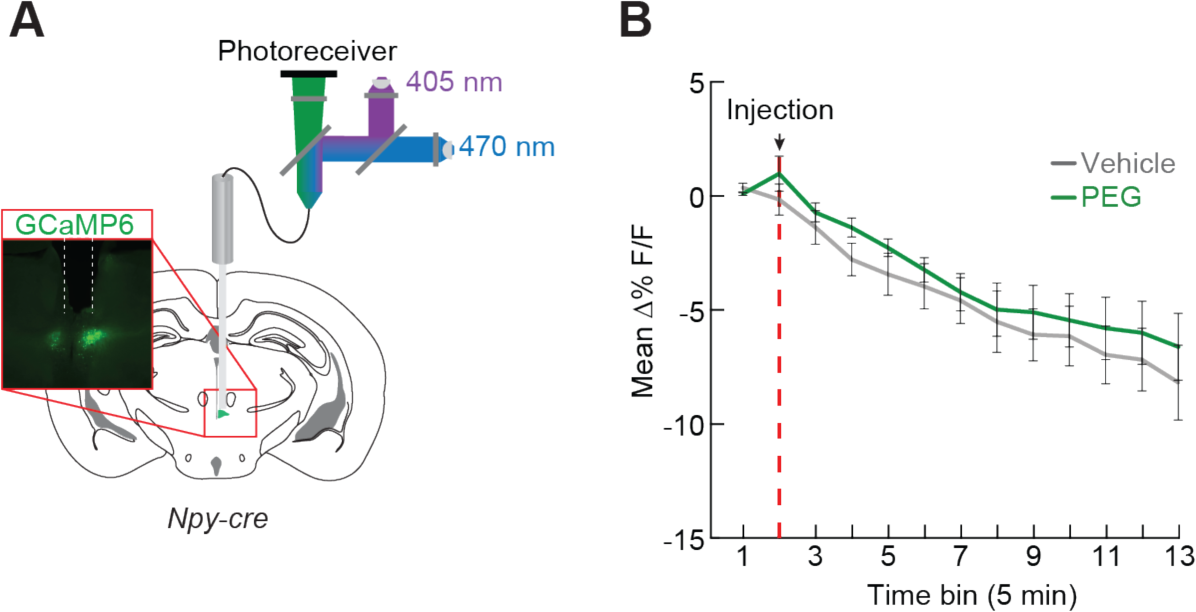
Subparafascicular NPY neurons are not activated by thirst. (**A**) Dual wavelength fiber photometry was used to record activity of subparafascicular nucleus NPY-expressing neurons. (**B**) Mean ΔF/F in 5 min bins of GCaMP6s signal after an injection of saline (grey) or PEG (green, n=5, repeated measure two-way ANOVA, n.s.). Data are expressed as mean ± SEM.

**Figure S7.**
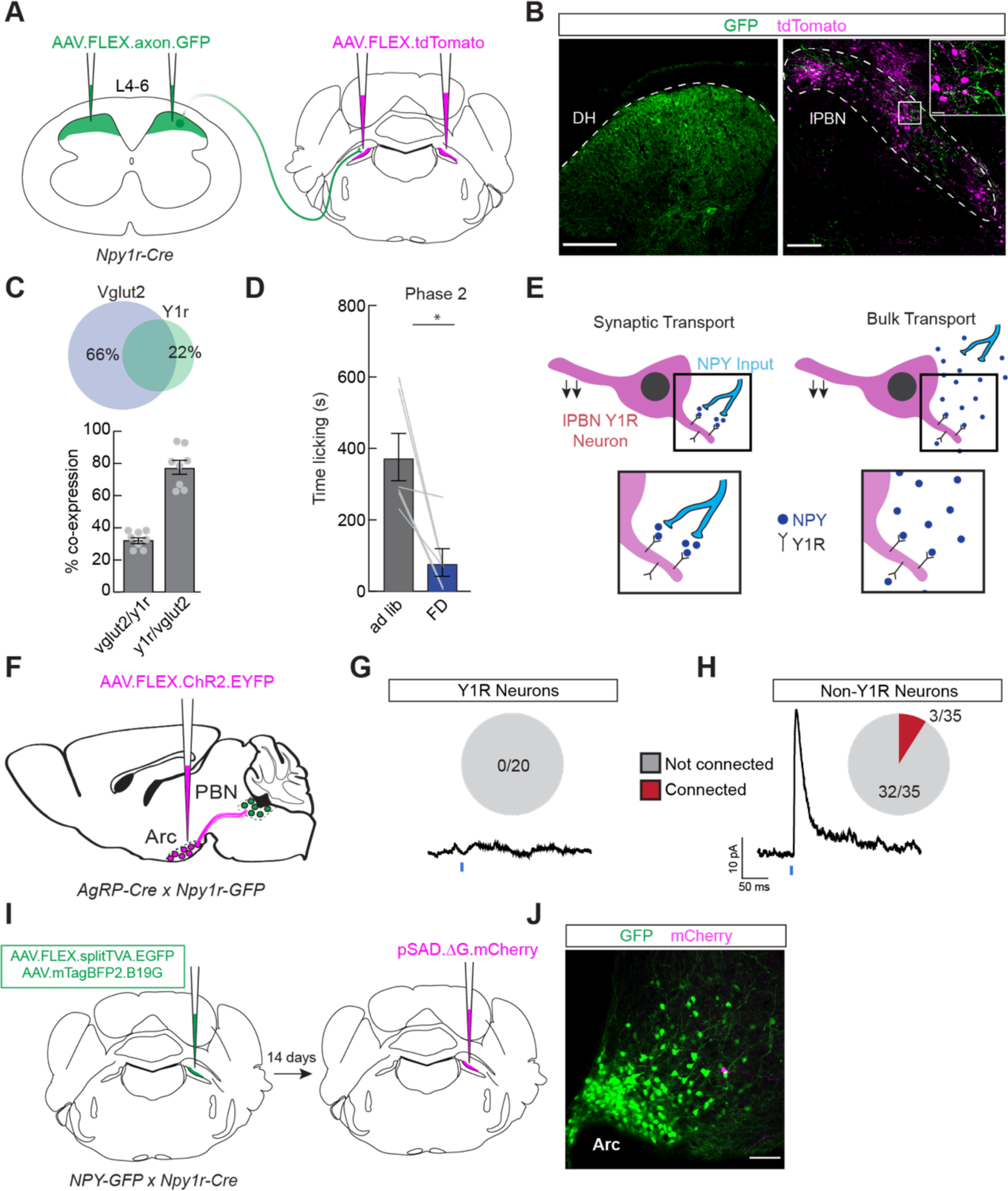
Glutamatergic lPBN Y1R neurons do not receive direct monosynaptic input from hypothalamic NPY expressing neurons. (**A**) Approach to visualize Y1R-expressing neurons in the lPBN and projections from the dorsal spinal cord. (**B**) Representative images showing Y1R neurons expressing GFP at the site of injection in the dorsal horn (left) and their projections to the lPBN along with Y1R-expressing lPBN neurons expressing tdTomato (right). Scale bar, 200 μm (inset, 10 μm). (**C**) (top) In situ hybridization results. Venn diagram depicting percent of cells expressing Vglut2 and Npy1r. Circle diameters are proportional to total number of cells expressing each gene. (bottom) Percent of cells co-expressing Vglut2 and Npy1r. Grey dots represent individual sections from 3 mice. (**D**) Time licking during Phase 2 in ad libitum fed (grey) or food deprived (blue) mice after genetic deletion of Y1R from dorsal horn neurons (n=6, paired t-test, p<0.05). (**E**) Two possible NPY signaling mechanisms: NPY inputs to the lPBN (activated by competing needs) making synaptic contact and acting on the Y1 receptor (left) or extra-synaptic diffusion of NPY onto Y1 receptors (right). (**F**) Channelrhodopsin-assisted circuit mapping – activation of arcuate NPY projections to the lPBN while monitoring activity in lPBN Y1R neurons. (**G**) Representative trace showing a lack of evoked current in an lPBN Y1R neuron following activation of NPY inputs (blue). None of the 20 patched neurons showed inhibitory post synaptic currents (IPSC, n=3 mice). (**H**) Representative trace of an evoked IPSC in a non-lPBN neuron following activation of NPY inputs (blue). 3/35 neurons patched showed inhibitory post synaptic potentials (n=3 mice). (**I**) Injection strategy for monosynaptic rabies tracing from lPBN Y1R neurons. EnvA-coated rabies virus (ι1G) was injected into the lPBN 14 days after helper injections. (**J**) Representative image of arcuate NPY neurons (green) and rabies expressing, presynaptic inputs (magenta). Scale bar, 200 μm;. Data are expressed as mean ± SEM. Grey lines represent individual mice. T-tests: *p<0.05.

**Figure S8.**
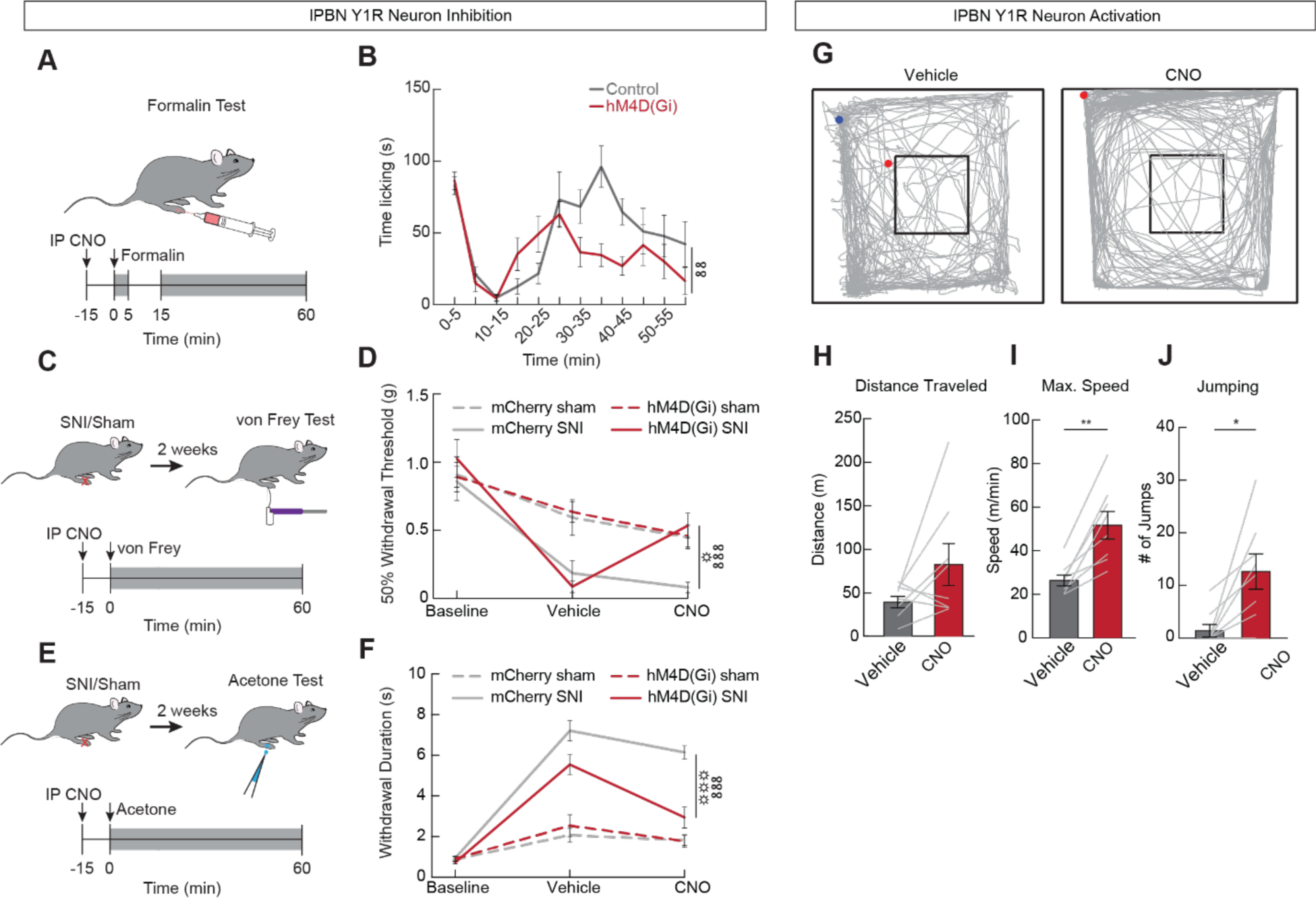
Inhibition of lPBN Y1R neurons suppresses sustained pain behaviors and activation evokes pain-like responses. (**A**) Experimental timeline of lPBN Y1R neuron inhibition during the formalin test. (**B**) Time spent licking paw in 5 min bins after an injection of formalin in mice injected with a control virus (grey) or mice injected with hM4D(Gi) to chemogenetically inhibit lPBN Y1R neurons (red, n=10-11, repeated measures two-way ANOVA, group x time interaction p<0.01). (**C**) Experimental timeline of lPBN Y1R neuron inhibition following SNI, von Frey experiments. (**D**) Withdrawal thresholds before SNI or sham (baseline), after SNI or sham in saline treated animals, and after SNI or sham in CNO treated animals. Grey lines represent mice injected with control virus, red lines represent mice injected with hM4D(Gi), and dotted lines represent sham animals (n=8/group, repeated measures three-way ANOVA, main effect of virus p<0.05). (**E**) Experimental timeline testing the effect of lPBN Y1R neuron inhibition following SNI, cold allodynia. (**F**) Withdrawal duration after application of acetone to the paw before SNI or sham (baseline), after SNI or sham in control saline treated animals, and after SNI or sham in CNO treated animals. Grey lines represent mice injected with control virus, red lines represent mice injected with hM4D(Gi), and dotted lines represent sham animals (n=8/group, repeated measures three-way ANOVA, main effect of virus p<0.001). (**G**) Representative plots showing locomotion of an hM3D(Gq)-expressing mouse in an open field after a control saline injection (left) or CNO induced activation of lPBN Y1R neurons (right). Blue dots represent the location of the mouse at the start of the assay and red dots represent the location at the end of the assay. (**H**) Distance traveled during the open field assay in control saline treated (grey) and CNO treated (red) mice (n=8, paired t-test, n.s.). (**I**) Maximum speed reached during the open field assay in control and CNO treated mice (n=8, paired t-test, p<0.01). (**J**) Number of jumps in hM3D(Gq)-expressing mice after a control saline (grey) or CNO (red) injection (n=8, paired t-test, p<0.05). Data are expressed as mean ± SEM. Grey lines represent individual mice. T-test and post-hoc comparisons: *p<0.05, **p<0.01. ANOVA main effect of group: ☼p<0.05, ☼☼☼p<0.001. ANOVA time x virus interaction: ∞∞p<0.01, ∞∞∞p<0.001.

**Figure S9.**
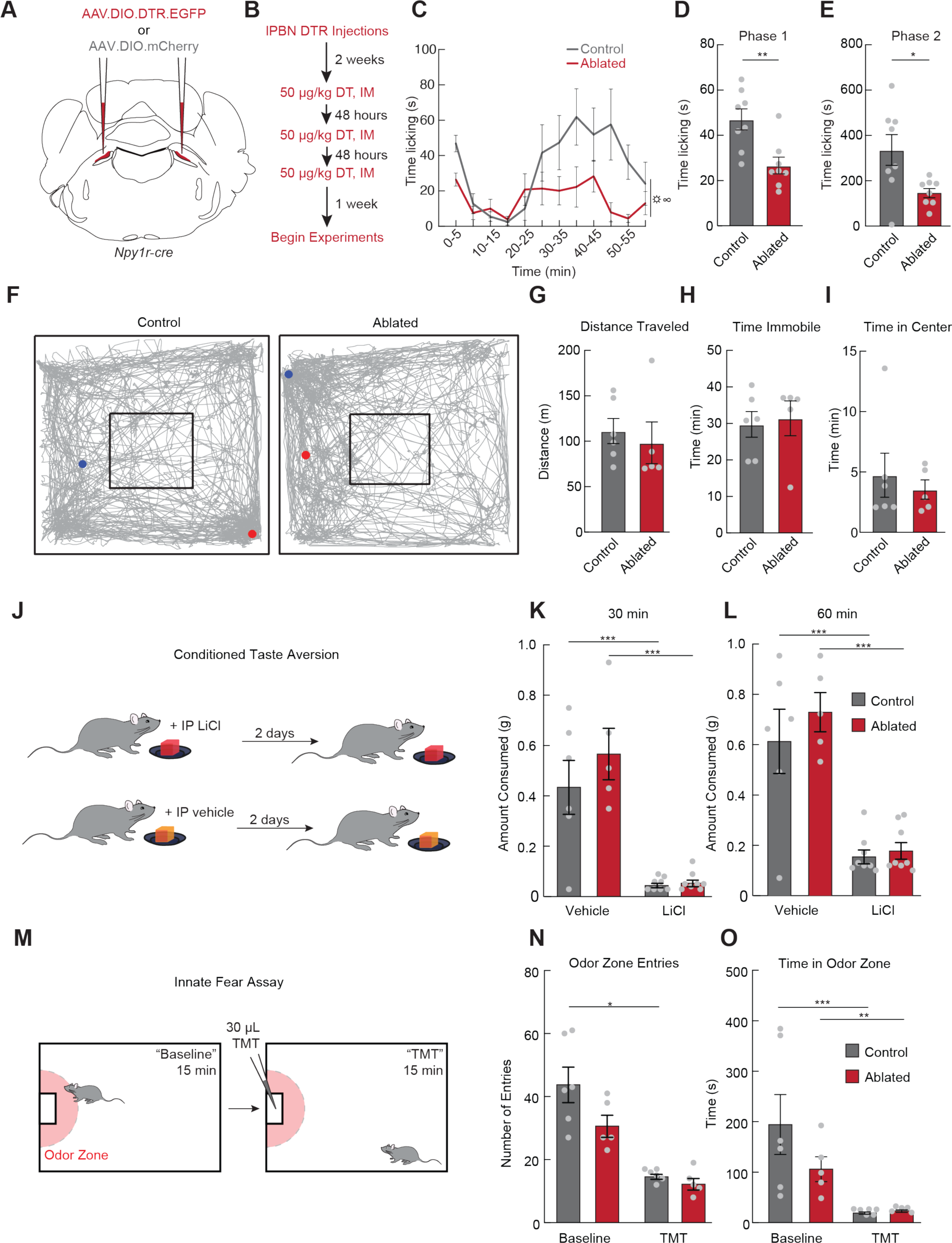
lPBN Y1R neuron ablation suppresses pain behaviors without affecting general locomotion or responses to other negative stimuli. (**A**) Injection strategy for bilateral lPBN Y1R neuron ablation. (**B**) Experimental timeline for lPBN Y1R neuron ablation. (**C**) Time spent licking paw in 5 min bins after an injection of formalin in control mice (grey) or Y1R ablated mice (red, n=8/group, repeated measures two-way ANOVA, main effect of group p<0.05). (**D**) Time spent licking paw during Phase 1 in (C) (n=8/group, unpaired t-test, p<0.01). (**E**) Time spent licking paw during Phase 2 in (C) (n=8/group, unpaired t-test, p<0.05). (**F**) Representative plots showing locomotion of control (left) and ablated (right) mice. Blue dots represent the location of the mouse at the start of the assay and red dots represent the location at the end of the assay. (**G-I**) Distance traveled (G), time spent immobile (H), and time spent in the center of the arena (I) in control (grey) and ablated (red) mice (n=5-6/group, unpaired t-tests, all n.s.). (**J**) Conditioned taste aversion paradigm. (**K-L**) Amount of gel paired with saline or LiCl consumed by control mice (grey) and Y1R ablated mice (red) after 30 min (K) and 60 mins (L) (n=5-8/group, two-way ANOVA, main effect of virus n.s.). (**M**) Predator odor avoidance paradigm. (**N**) Number of entries into the odor zone before and after addition of TMT in control mice (grey) and Y1R ablated mice (red, n=5-6/group, repeated measures two-way ANOVA, main effect of virus n.s.). (**O**) Time spent in the odor zone before and after addition of TMT in control mice (grey) and Y1R ablated mice (red, n=5-6/group, repeated measures two-way ANOVA, main effect of virus n.s.). Data are expressed as mean ± SEM. Grey dots represent individual mice. T-test and post-hoc comparisons: *p<0.05, **p<0.01, ***p<0.001. ANOVA main effect of group: ☼p<0.05. ANOVA interaction: ∞p<0.05.

**Figure S10.**
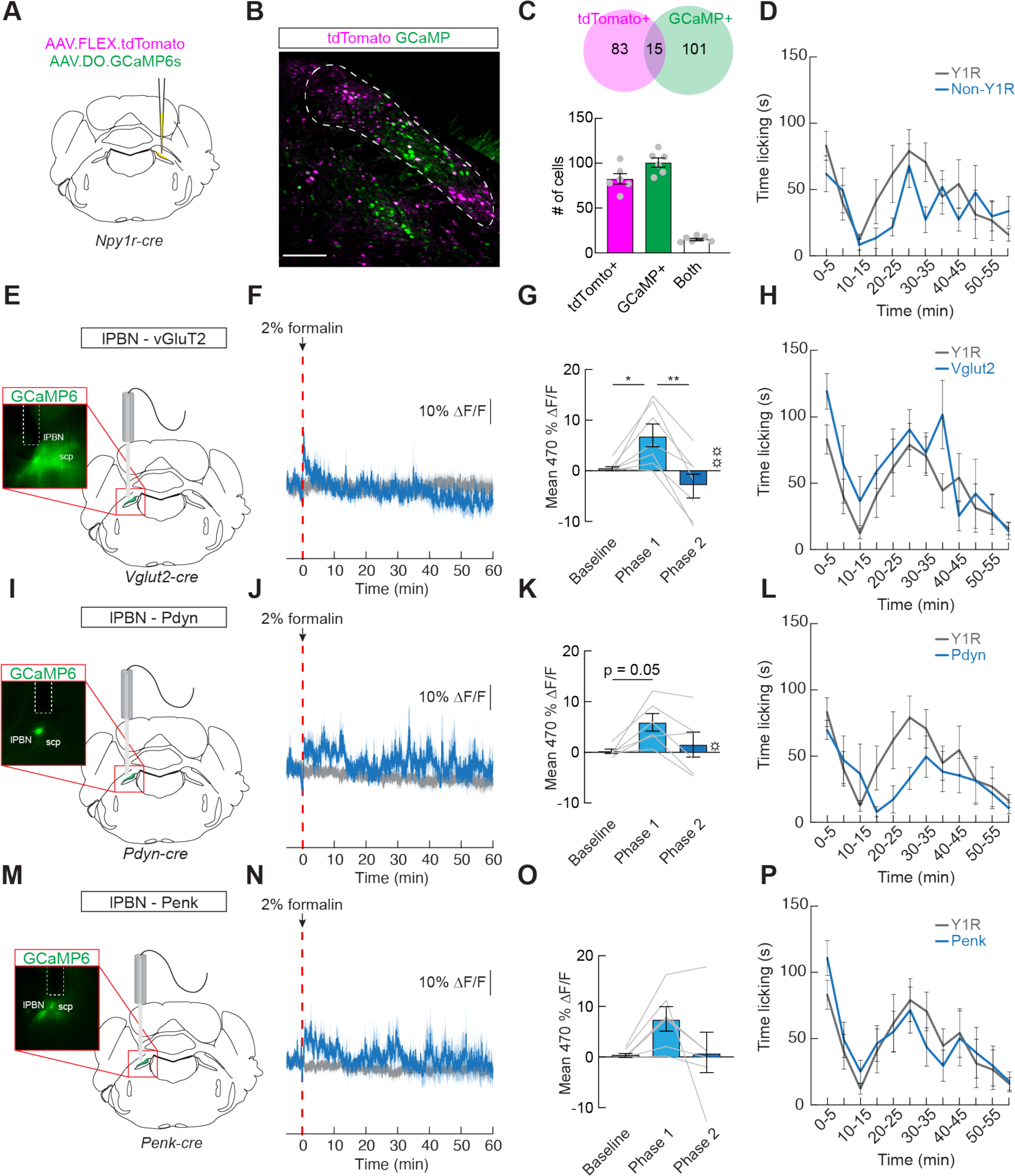
Activity of non-Y1R lPBN cell types does not show sustained, slow components that may code for pain state. (**A**) Injection strategy to confirm specificity of Cre-OFF GCaMP expression in an *Npy1r:Cre* mouse. (**B**) Representative image showing Cre-OFF GCaMP (green) and Cre-dependent tdTomato (red) expression in the PBN. Scale bar, 200 μm. (**C**) (top) Number of cells expressing only tdTomato (red), only GCaMP (green), or both (yellow) in the PBN. (bottom) Average number of neurons expressing tdTomato, GCaMP, and both. Grey dots represent individual sections. (**D**) Time spent licking paw in 5 min bins after an injection of formalin while recording from Y1R neurons (grey) or non-Y1R neurons (blue, n=6-9/group, repeated measures two-way ANOVA, n.s.) (**E**) Fiber photometry was used to measure calcium dynamics of lPBN Vglut2-expressing neurons in awake, behaving mice. Inset, representative image showing GCaMP6s expression in lPBN Vglut2 neurons below a fiber optic tract. (**F**) Average ι1F/F of GCaMP6s signal from lPBN Vglut2 neurons after a formalin injection into the dorsal surface of the hindpaw (dotted red line). Signals are aligned to formalin injection. Blue, 470 nm; grey, 405 nm. Dark lines represent mean and lighter, shaded areas represent SEM. (**G**) Mean ι1F/F of lPBN Vglut2 neurons before formalin injection, during Phase 1, and during Phase 2 (n=7, repeated measures one-way ANOVA, p<0.01). (**H**) Time spent licking paw in 5 min bins after an injection of formalin while recording from Y1R neurons (grey) or Vglut2 neurons (blue, n=7-9/group, repeated measures two-way ANOVA, n.s.). (**I**) Fiber photometry was used to measure calcium dynamics of lPBN Pdyn-expressing neurons. Inset, representative image showing GCaMP6s expression in lPBN Pdyn neurons below a fiber optic tract. (**J**) Average ι1F/F of GCaMP6s signal from lPBN Pdyn neurons after a formalin injection. (**K**) Mean ι1F/F of lPBN Pdyn neurons before formalin injection, during Phase 1, and during Phase 2 (n=6, repeated measures one-way ANOVA, main effect of time p<0.05). (**L**) Time spent licking in 5 min bins after an injection of formalin while recording from Y1R neurons (grey) or Pdyn neurons (blue, n=6-9/group, repeated measures two-way ANOVA, n.s.). (**M**) Fiber photometry was used to measure calcium dynamics of lPBN Penk-expressing neurons. Inset, representative image showing GCaMP6s expression in lPBN Penk neurons below a fiber optic tract. (**N**) Average ι1F/F of GCaMP6s signal from lPBN Penk neurons after a formalin injection. (**O**) Mean ι1F/F of lPBN Penk neurons before formalin injection, during Phase 1, and during Phase 2 (n=6, repeated measures one-way ANOVA, n.s.). (**P**) Time spent licking in 5 min bins after an injection of formalin while recording from Y1R neurons (grey) or Penk neurons (blue, n=6-9/group, repeated measures two-way ANOVA, n.s.). Data are expressed as mean ± SEM. Grey lines represent individual mice. ANOVA main effect of group: ☼p<0.05, ☼☼p<0.01. Post-hoc comparisons: *p<0.05. **p<0.01.

**Figure S11.**
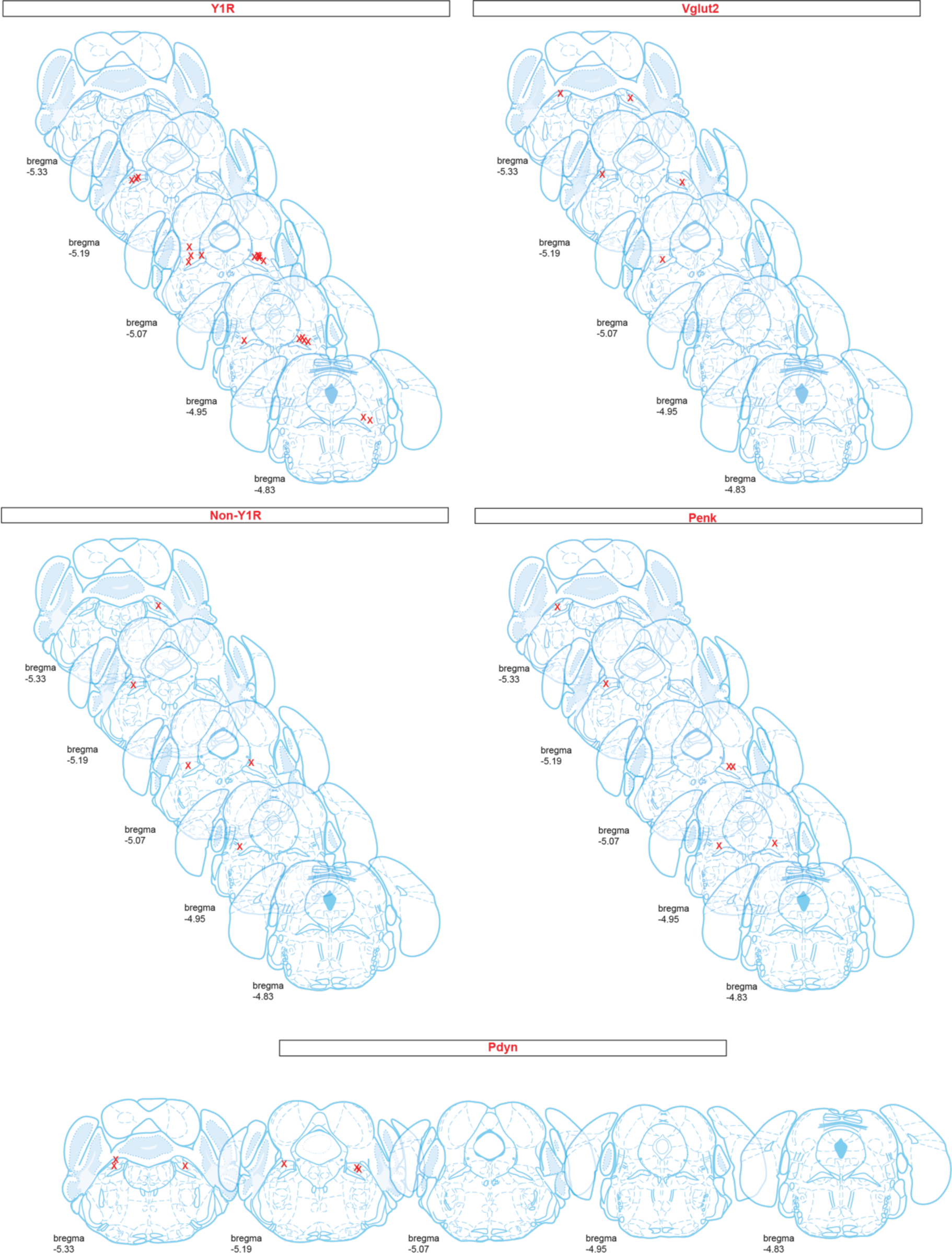
Fiber placement from fiber photometry experiments. Diagrams depicting fiber tip location determined from histological examination of fiber photometry brains from all genotypes used.

**Figure S12.**
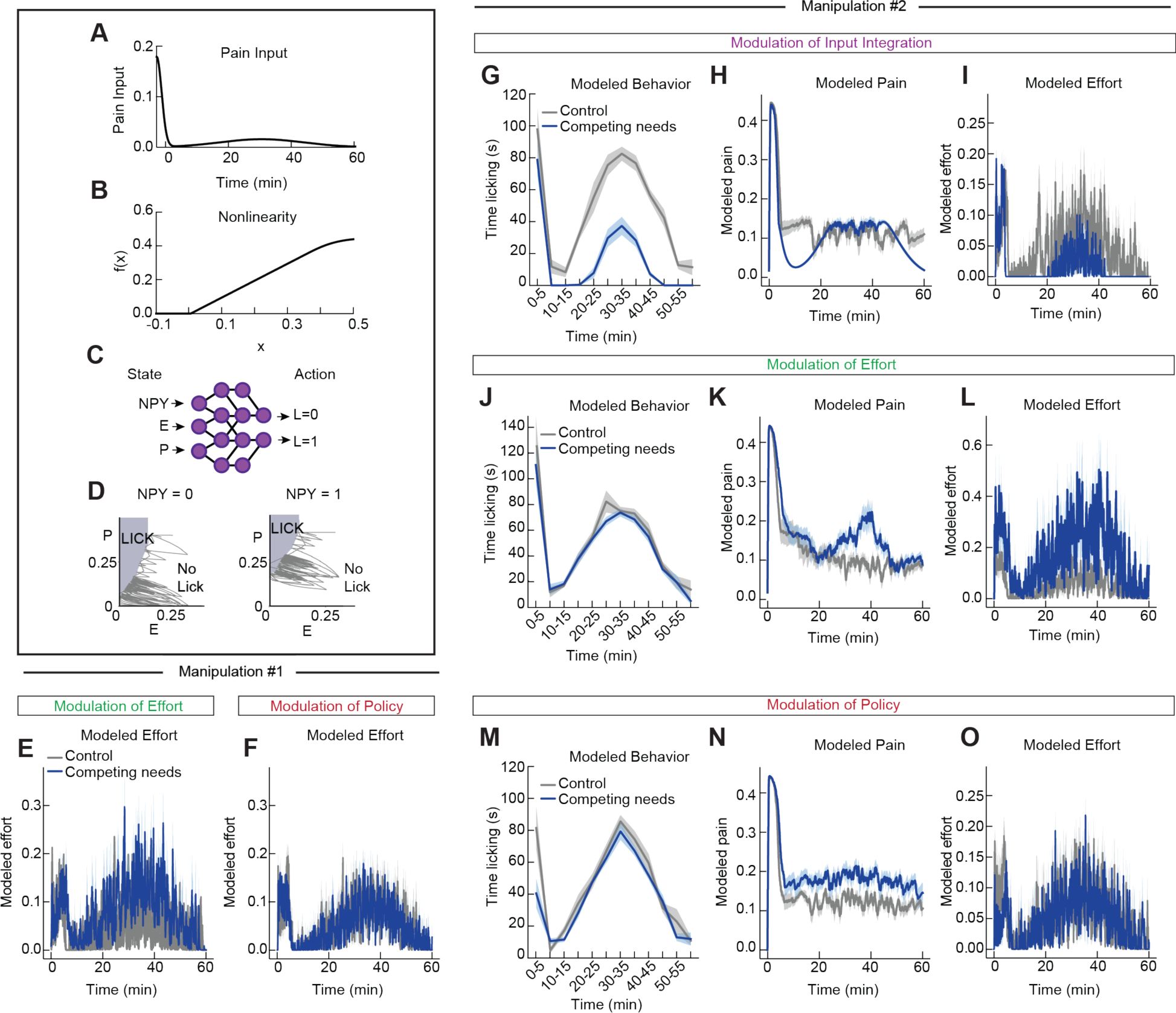
Modeling pain behavior, pain state, and effort following formalin administration. (**A**) The average pain input given when training and simulating the model, chosen as the sum of two Gaussians corresponding to phases 1 and 2 of the formalin response. Model input during training and simulation is a noisy version of this input. (**B**) The pain state of the model is an integral of the pain input passed through a rectified saturating nonlinearity, shown here. (**C**) Schematic of the neural network implementing the behavioral policy, also showing the addition of a third state dimension ‘NPY’ that is 1 in the presence of a competing survival need and 0 otherwise (used in “Modulation of Policy” simulations). (**D**) Example policy space after training one model in the “Modulation of Policy” scenario in which the model learns a distinct policy for the presence vs absence of a competing need state. The dark gray line indicates a trajectory in the pain-effort state space, while the light gray region indicates the portion of pain-effort state space in which the model produces a lick response (no lick response is produced outside this region.) When NPY = 1, the lick region shifts upwards compared to NPY = 0, corresponding to an increase in the pain threshold required to produce a licking response. (**E,F**) Average ‘effort’ dynamics of 8 trained models for the baseline model (‘control’) vs a model that increases the effort cost of licking (E) or introduces a third axis to the behavioral control policy (F). Dark lines represent mean and lighter, shaded areas represent SEM. (**G-O**) Simulation results using alternative formulations of input integration, effort modulation, and policy modulation (see equation 2 for each manipulation in the Methods.) (**G-I**) Simulated licking behavior (G), ‘pain’ state (H) and ‘effort’ state (I) in the baseline condition (‘control’) vs a competing need that alters integration of nociceptive input using manipulation 2. (**J-L**) Simulated licking behavior (J), ‘pain’ state (K) and ‘effort’ state (L) in the baseline condition (‘control’) vs a competing need that alters the effort cost of licking using manipulation 2. (**M-O**) Simulated licking behavior (M), ‘pain’ state (N) and ‘effort’ state (O) in the baseline condition (‘control’) vs a competing need that adds a third dimension to the behavioral control policy using manipulation 2. Data are expressed as mean ± SEM.

**Figure S13.**
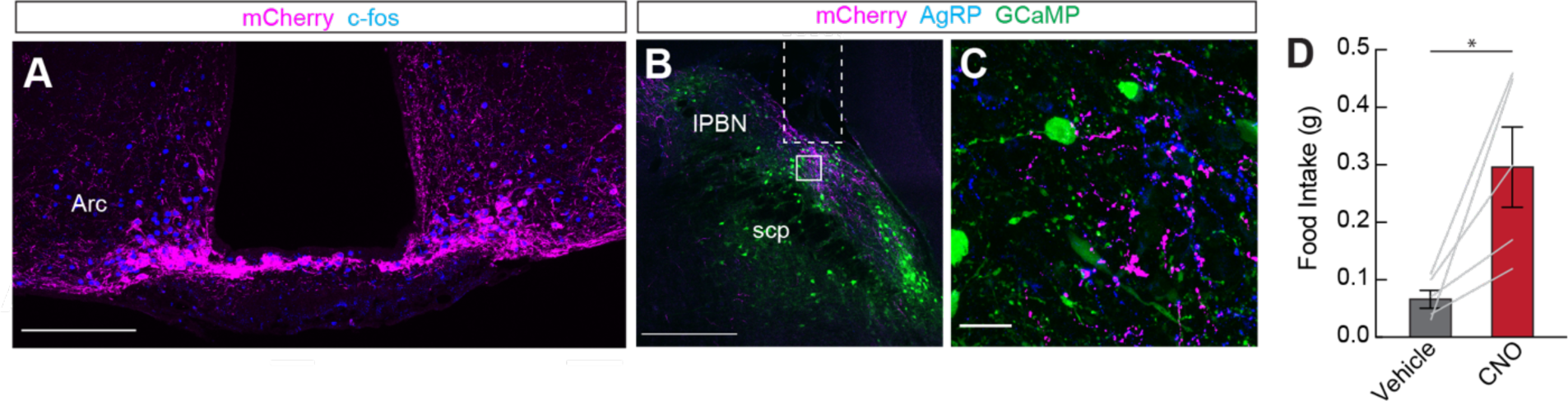
Validation of DREADD expression in Y1R photometry mice. (**A**) Representative image showing hM3D(Gq)-mCherry expression in the arcuate nucleus with c-fos expression confirming neural activation after a CNO injection. Scale bar, 200 μm. (**B**) Representative image of the lPBN showing GCaMP expression in Y1R neurons and hM3(Gq)-mCherry-expressing axons of arcuate neurons. The fiber tract is also visible (dotted white). Scale bar, 200 μm. (**C**) High resolution image of the area inside the white square show in (B). Scale bar, 20 μm. (**D**) Food consumed after an injection of saline (grey) or CNO (red, n=5, paired t-test, p<0.05). Data are expressed as mean ± SEM. Grey lines represent individual mice. T-test: *p<0.05.

**Figure S14.**
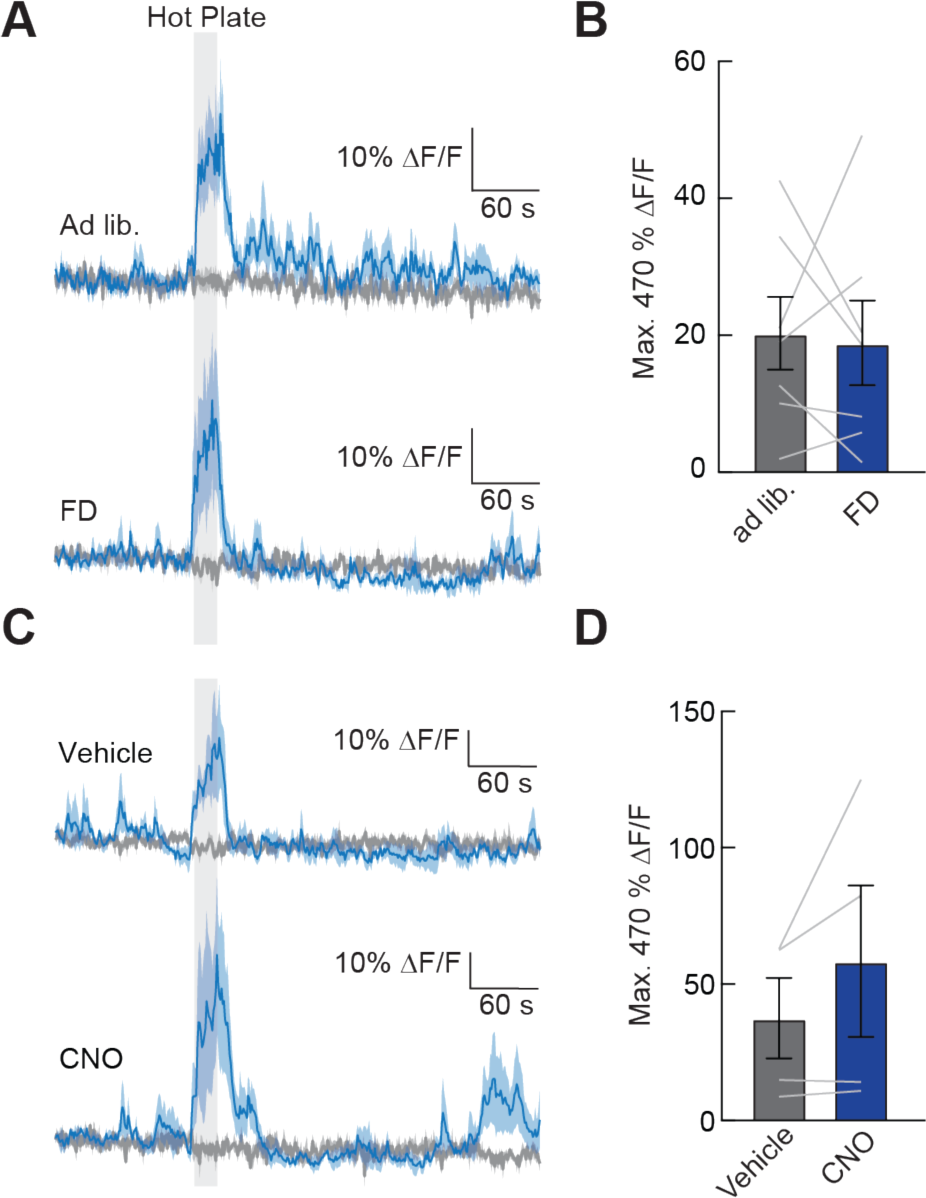
Food deprivation and arcuate NPY neuron activation does not affect lPBN Y1R neuron responses to acute noxious heat. (**A**) Average ΔF/F of GCaMP6s signal from lPBN Y1R neurons while *ad libitum* fed or food deprived mice were placed on a 52° C hot plate for 20 s (grey box). Signals are aligned to hot plate onset. Blue, 470 nm; grey, 405 nm. Dark lines represent mean and lighter, shaded areas represent SEM. (**B**) Maximum ΔF/F of GCaMP6s signals shown in (A) during hot plate period shown in in grey (n=7, paired t-test, n.s.) (**C**) Average ΔF/F of GCaMP6s signal during hot plate assay from lPBN Y1R neurons in mice expressing hM3D(Gq) in arcuate NPY neurons after injection of saline control or CNO. (**D**) Maximum ΔF/F of GCaMP6s signals shown in (C) during time on the hot plate depicted by grey shading (n=4, paired t-test, n.s.). Data are expressed as mean ± SEM. Grey lines represent individual mice.

**Table S1.** Sample sizes, statistical tests, and results used in main figures.

| Figure | Sample Size | Statistical Test | Values |
| --- | --- | --- | --- |
| 1E | N = 7-9/group | Unpaired t-test | **p=0.0025 |
| 1F | N = 15 | RM one-way ANOVA<br>Post hoc comparison: Holm-Sidak<br>Baseline vs. CFA – ad lib<br>Baseline vs. CFA – FD<br>CFA – ad lib vs. CFA – FD | F(1.66,23.19)=48.35, ***p<0.001<br>***p<0.001<br>**p=0.0058<br>***p<0.001 |
| 1G | N = 8 | RM one-way ANOVA<br>Post hoc comparison: Holm-Sidak<br>Baseline vs. SNI – ad lib<br>Baseline vs. SNI – FD<br>SNI – ad lib vs. SNI – FD | F(1.51,10.55)=33.27, ***P<0.001<br>***p<0.001<br>*p=0.0180<br>**p=0.0025 |
| 1I | N = 9/group | Unpaired t-test | **p=0.0043 |
| 1J | N = 8 | RM one-way ANOVA<br>Post hoc comparison: Holm-Sidak<br>Baseline vs. CFA – Veh<br>Baseline vs. CFA – PEG<br>CFA – Veh vs. CFA – PEG | F(1.49,10.42)=8.704, **p=0.0088<br>*p=0.0213<br>p=0.1691<br>*p=0.0213 |
| 1K | N = 10 | RM one-way ANOVA<br>Post hoc comparison: Holm-Sidak<br>Baseline vs. SNI – Veh<br>Baseline vs. SNI – PEG<br>SNI – Veh vs. SNI – PEG | F(1.50,13.54)=296.0, ***p<0.001<br>***p<0.001<br>***p<0.001<br>***p<0.001 |
| 1M | N = 8/group | Unpaired t-test | **p=0.0012 |
| 1O | N = 10-12/group | Unpaired t-test | *p=0.0495 |
| 2B | N = 7-8/group | RM two-way ANOVA<br>Factor 1: group (infusion)<br>Factor 2: time<br>Interaction: group x time<br>Post hoc comparison: Holm-Sidak<br>0-5 min: VEH vs. Antag.<br>5-10 min: VEH vs. Antag.<br>10-15 min: VEH vs. Antag.<br>15-20 min: VEH vs. Antag.<br>20-25 min: VEH vs. Antag.<br>25-30 min: VEH vs. Antag.<br>30-35 min: VEH vs. Antag.<br>35-40 min: VEH vs. Antag.<br>40-45 min: VEH vs. Antag.<br>45-50 min: VEH vs. Antag.<br>50-55 min: VEH vs. Antag.<br>55-60 min: VEH vs. Antag. | F(1,13)=6.305, *p=0.0260<br>F(3.12, 40.52)=3.838, *p=0.0155<br>F(11, 143)=0.6946, p=0.7421<br>p=0.9789<br>p=0.9563<br>p=0.9563<br>p=0.8274<br>p=0.8222<br>p=0.8679<br>p=0.8267<br>p=0.9789<br>p=0.9563<br>p=0.9563<br>p=0.8267<br>p=0.9563 |
| 2C | N = 7-8/group | Unpaired t-test | p=0.6295 |
| 2D | N = 7-8/group | Unpaired t-test | *p=0.0137 |
| 2I | N = 5 | RM two-way ANOVA<br>Factor 1: group (odor)<br>Factor 2: time<br>Interaction: group x time<br>Post hoc comparison: Holm-Sidak<br>-5 min – -4 min: VEH vs. TMT<br>-4 min – -3 min: VEH vs. TMT<br>-3 min – -2 min: VEH vs. TMT<br>-2 min – -1 min: VEH vs. TMT<br>-1 min – 0 min: VEH vs. TMT<br>0 min – 1 min: VEH vs. TMT<br>1 min – 2 min: VEH vs. TMT | F(1,8)=5.033, p=0.0551<br>F(1.36,10.87)=1.50, p=0.2572<br>F(9,72)=2.239, *p=0.0288<br>p=0.9119<br>p=0.9099<br>p=0.9439<br>p=0.9119<br>p=0.8523<br>p=0.8266<br>p=0.6809 |
|  |  | 2 min – 3 min: VEH vs. TMT<br>3 min – 4 min: VEH vs. TMT<br>4 min – 5 min: VEH vs. TMT | p=0.3189<br>p=0.5091<br>p=0.5091 |
| 2K | N = 6 | RM two-way ANOVA<br>Factor 1: group<br>Factor 2: time<br>Interaction: group x time<br>Post hoc comparison: Holm-Sidak<br>0-5 min: ad lib. vs. FD<br>5-10 min: ad lib. vs. FD<br>10-15 min: ad lib. vs. FD<br>15-20 min: ad lib. vs. FD<br>20-25 min: ad lib. vs. FD<br>25-30 min: ad lib. vs. FD<br>30-35 min: ad lib. vs. FD<br>35-40 min: ad lib. vs. FD<br>40-45 min: ad lib. vs. FD<br>45-50 min: ad lib. vs. FD<br>50-55 min: ad lib. vs. FD<br>55-60 min: ad lib. vs. FD | F(1,5)=13.47, *p=0.0144<br>F(2.55,12.77)=3.241, p=0.0636<br>F(2.5,12.51)=1.275, p=0.3206<br>p=0.8446<br>p=0.9677<br>p=0.7388<br>p=0.8446<br>p=0.9677<br>p=0.7656<br>p=0.7187<br>p=0.2550<br>p=0.1286<br>p=0.2722<br>p=0.7388<br>p=0.7656 |
| 3B | N = 10-11/group | Unpaired t-test | *p=0.0352 |
| 3C | N = 8 | RM one-way ANOVA<br>Post hoc comparison: Holm-Sidak<br>Baseline vs. SNI – Veh<br>Baseline vs. SNI – CNO<br>SNI – Veh vs. SNI – CNO | F(1.33, 9.29)=31.41, ***p<0.001<br>***p<0.001<br>*p=0.0133<br>***p<0.001 |
| 3D | N = 10-11/group | Unpaired t-test | p=0.6920 |
| 3E | N = 14 | Paired t-test | p=0.9611 |
| 3H | N = 6-7/group | RM two-way ANOVA<br>Factor 1: group (virus)<br>Factor 2: time (BL vs. test)<br>Interaction: group x time<br>Post hoc comparison: Holm-Sidak<br>Baseline: control vs. hM4D(Gi)<br>Test: control vs. hM4D(Gi) | F(1,11)=29.52, ***p<0.001<br>F(1,11)=51.70, ***p<0.001<br>F(1,11)=34.68, ***p<0.001<br>p=0.9211<br>***p<0.001 |
| 3I | N = 8 | Paired t-test | p=0.0546 |
| 4C | N = 9 | RM one-way ANOVA<br>Post hoc comparison: Holm-Sidak<br>Baseline vs. Phase 1<br>Baseline vs. Phase 2<br>Phase 1 vs. Phase 2 | F(1.64,13.10)=27.1, ***p<0.001<br>***p<0.001<br>*p=0.0111<br>**p=0.0032 |
| 4F | N = 9 | Paired t-test (on Fisher's z values) | **p=0.0055 |
| 4I | N = 6 | RM one-way ANOVA<br>Post hoc comparison: Holm-Sidak<br>Baseline vs. Phase 1<br>Baseline vs. Phase 2<br>Phase 1 vs. Phase 2 | F(1.09, 5.46)=3.365, p=0.1204<br>p=0.3160<br>p=0.7720<br>p=0.3160 |
| 4L | N = 6-9/group | Unpaired t-test (on Fisher's z values) | ***p<0.001 |
| 5O | N = 7 | RM one-way ANOVA<br>Post hoc comparison: Holm-Sidak<br>Baseline vs. Phase 1<br>Baseline vs. Phase 2<br>Phase 1 vs. Phase 2 | F(1.47, 8.80)=7.732, *p=0.0156<br>*p=0.0205<br>p=0.09384<br>p=0.0684 |
| 5Q | N = 7 | RM one-way ANOVA<br>Post hoc comparison: Holm-Sidak | F(1.15, 6.90)=3.818, p=0.0895 |
|  |  | Baseline vs. Phase 1<br>Baseline vs. Phase 2<br>Phase 1 vs. Phase 2 | p=0.2128<br>p=0.7336<br>p=0.2128 |
| 5S | N = 4 | RM one-way ANOVA<br>Post hoc comparison: Holm-Sidak<br>Baseline vs. Phase 1<br>Baseline vs. Phase 2<br>Phase 1 vs. Phase 2 | F(1.40,4.20)=8.829, *p=0.0348<br>p=0.1423<br>p=0.6819<br>p=0.1423 |
| 5T | N = 4-9/group | One-way ANOVA (on fisher's z values) | F(3,23)=0.1312, p=0.9405 |
| 5W | N = 5 | RM one-way ANOVA<br>Post hoc comparison: Holm-Sidak<br>Baseline vs. CFA – ad lib<br>Baseline vs. CFA – FD<br>CFA – ad lib vs. CFA – FD | F(1.36, 5.46)=10.82, *p=0.0158<br>**p=0.0099<br>p=0.4202<br>*p=0.0435 |
| 5X | N = 5 | RM two-way ANOVA<br>Factor 1: group<br>Factor 2: time<br>Interaction: group x time<br>Post hoc comparison: Holm-Sidak<br>-60 - -40s: BL vs. CFA – ad lib<br>-60 - -40s: BL vs. CFA – FD<br>-60 - -40s: CFA – ad lib v. CFA – FD<br>-40 - -20s: BL vs. CFA – ad lib<br>-40 - -20s: BL vs. CFA – FD<br>-40 - -20s: CFA – ad lib v. CFA – FD<br>-20 - 0s: BL vs. CFA – ad lib<br>-20 - 0s: BL vs. CFA – FD<br>-20 - 0s: CFA – ad lib v. CFA – FD<br>0 - 20s: BL vs. CFA – ad lib<br>0 - 20s: BL vs. CFA – FD<br>0 - 20s: CFA – ad lib v. CFA – FD<br>20 - 40s: BL vs. CFA – ad lib<br>20 - 40s: BL vs. CFA – FD<br>20 - 40s: CFA – ad lib v. CFA – FD<br>40 - 60s: BL vs. CFA – ad lib<br>40 - 60s: BL vs. CFA – FD<br>40 - 60s: CFA – ad lib v. CFA – FD | F(1.24,4.96)=5.926, p=0.0560<br>F(1.61,6.45)=26.80, ***p<0.001<br>F(2.40,9.61)=5.486, *p=0.0221<br>p=0.6307<br>p=0.6307<br>p=0.1103<br>p=0.0592<br>p=0.4006<br>p=0.4046<br>p=0.8758<br>p=0.4579<br>p=0.8856<br>*p=0.0165<br>p=0.0694<br>p=0.3042<br>**p=0.0044<br>p=0.2413<br>p=0.0563<br>p=0.4294<br>p=0.5815<br>p=0.3944 |

**Table S2.**
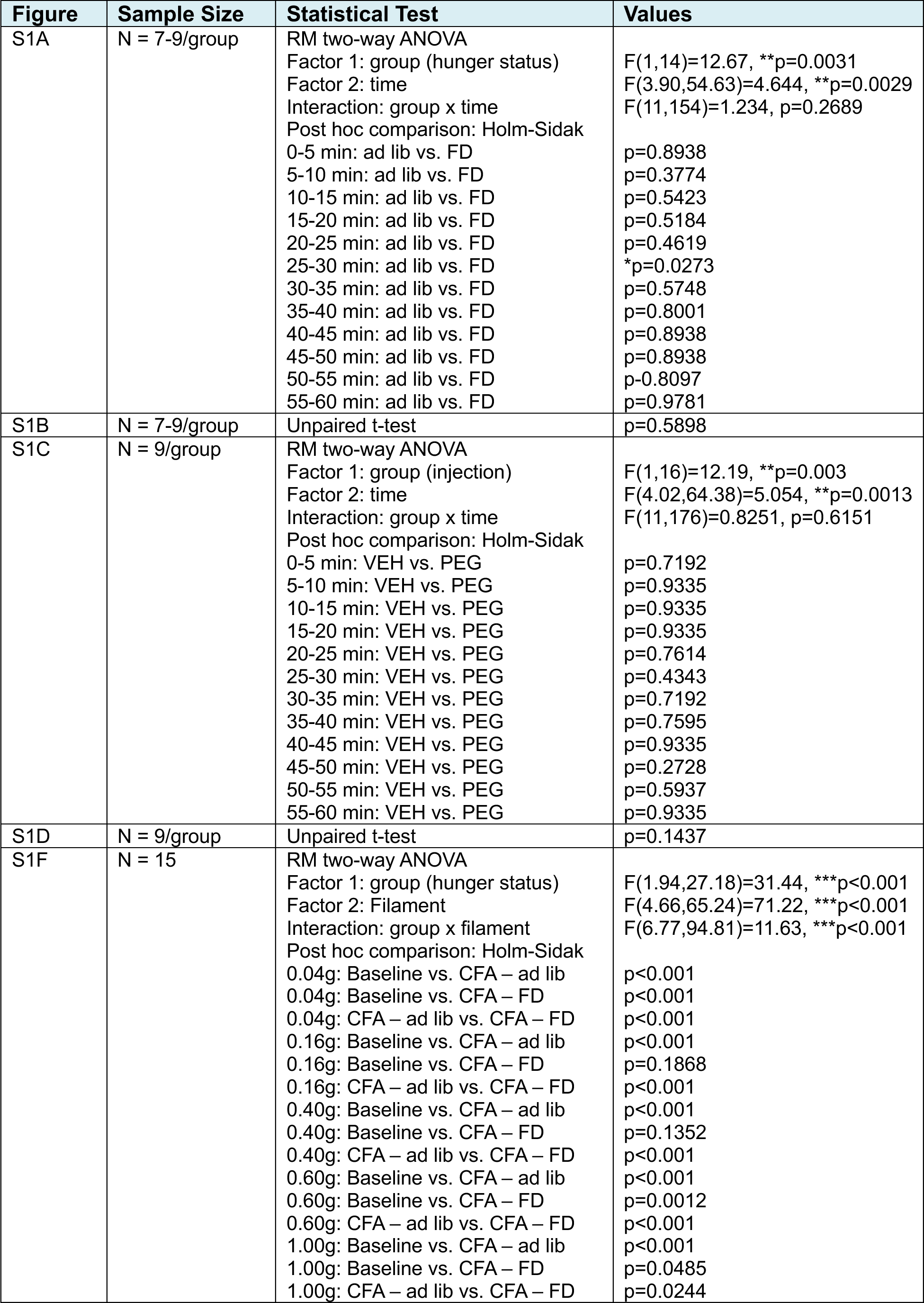
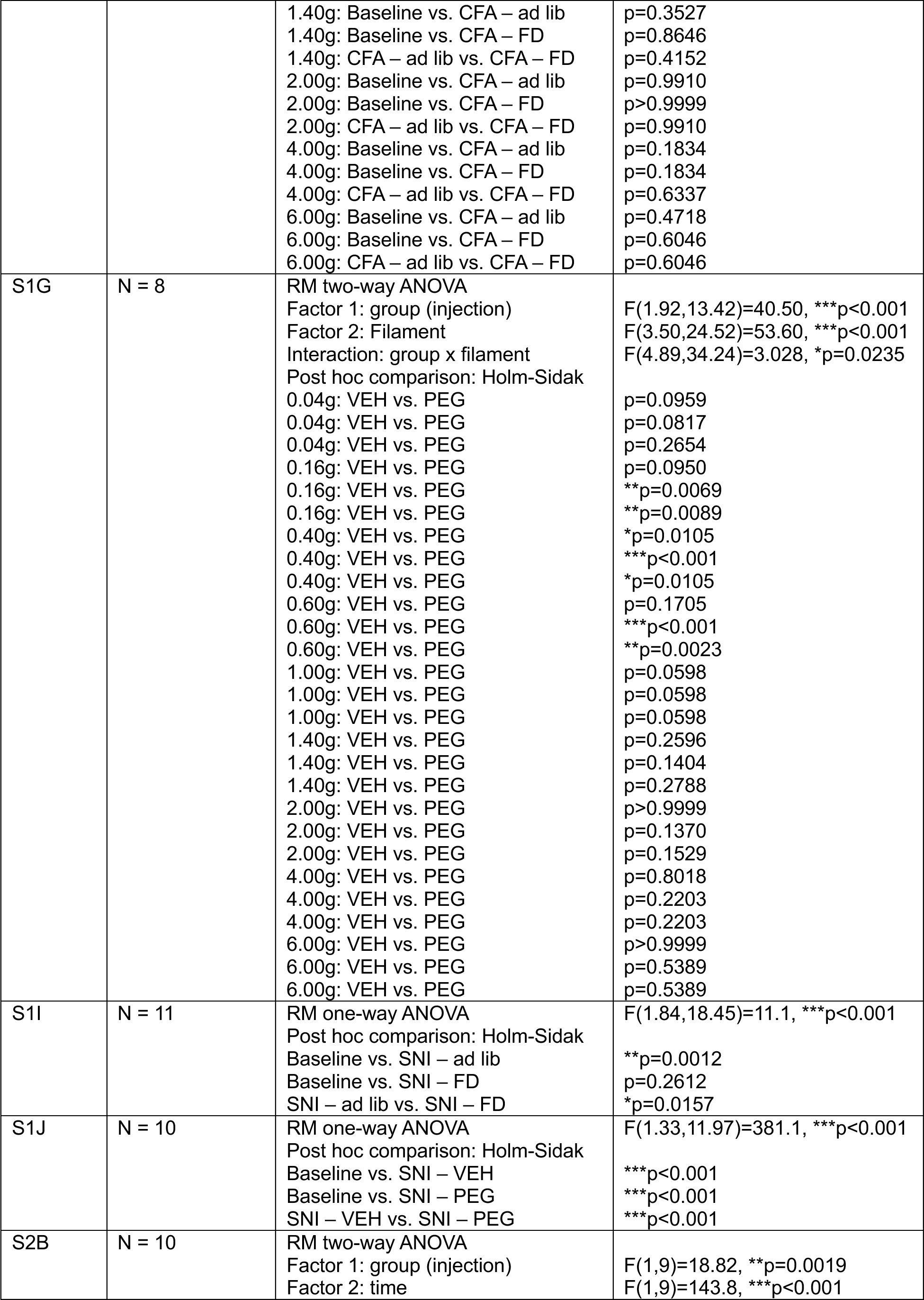
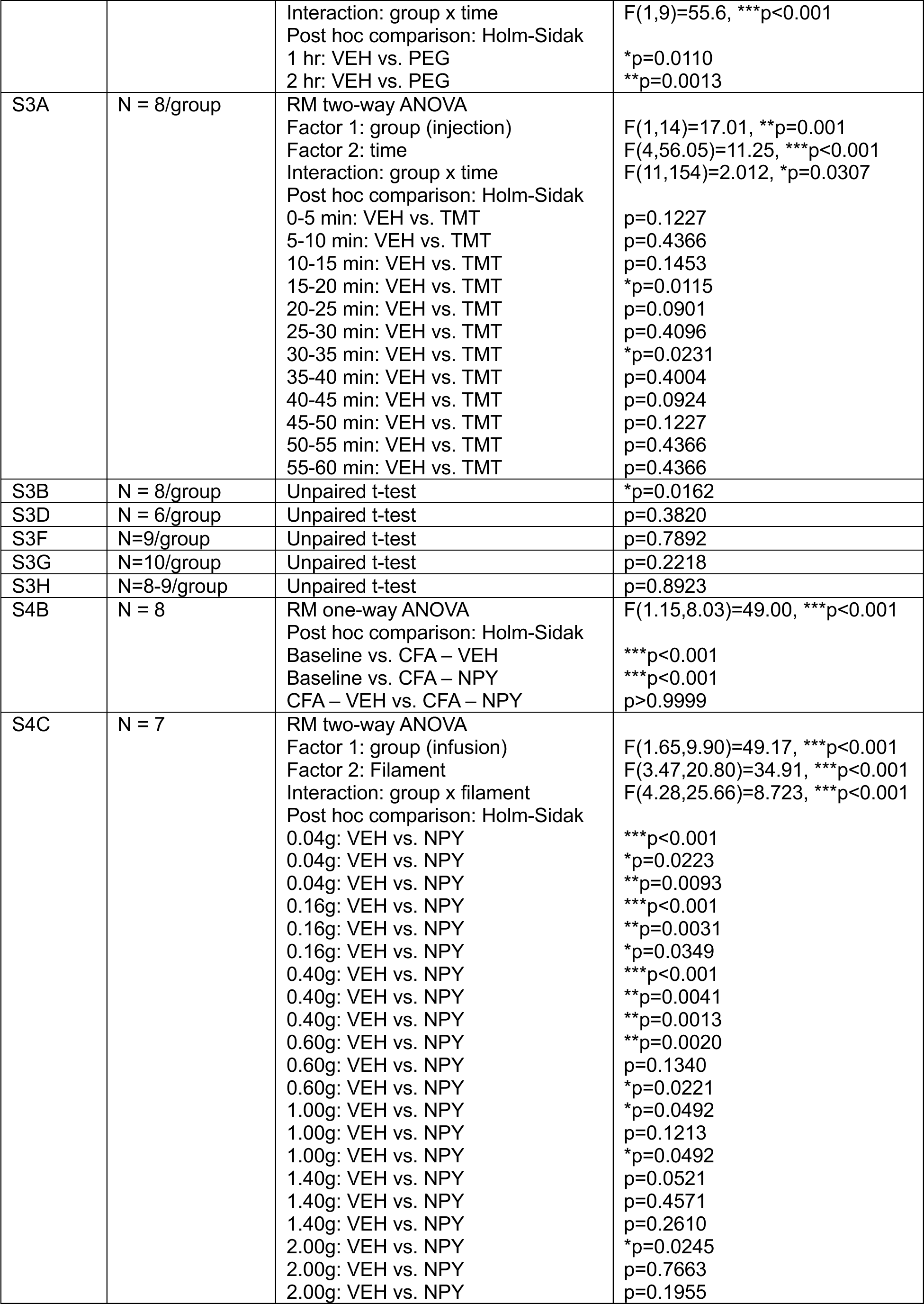
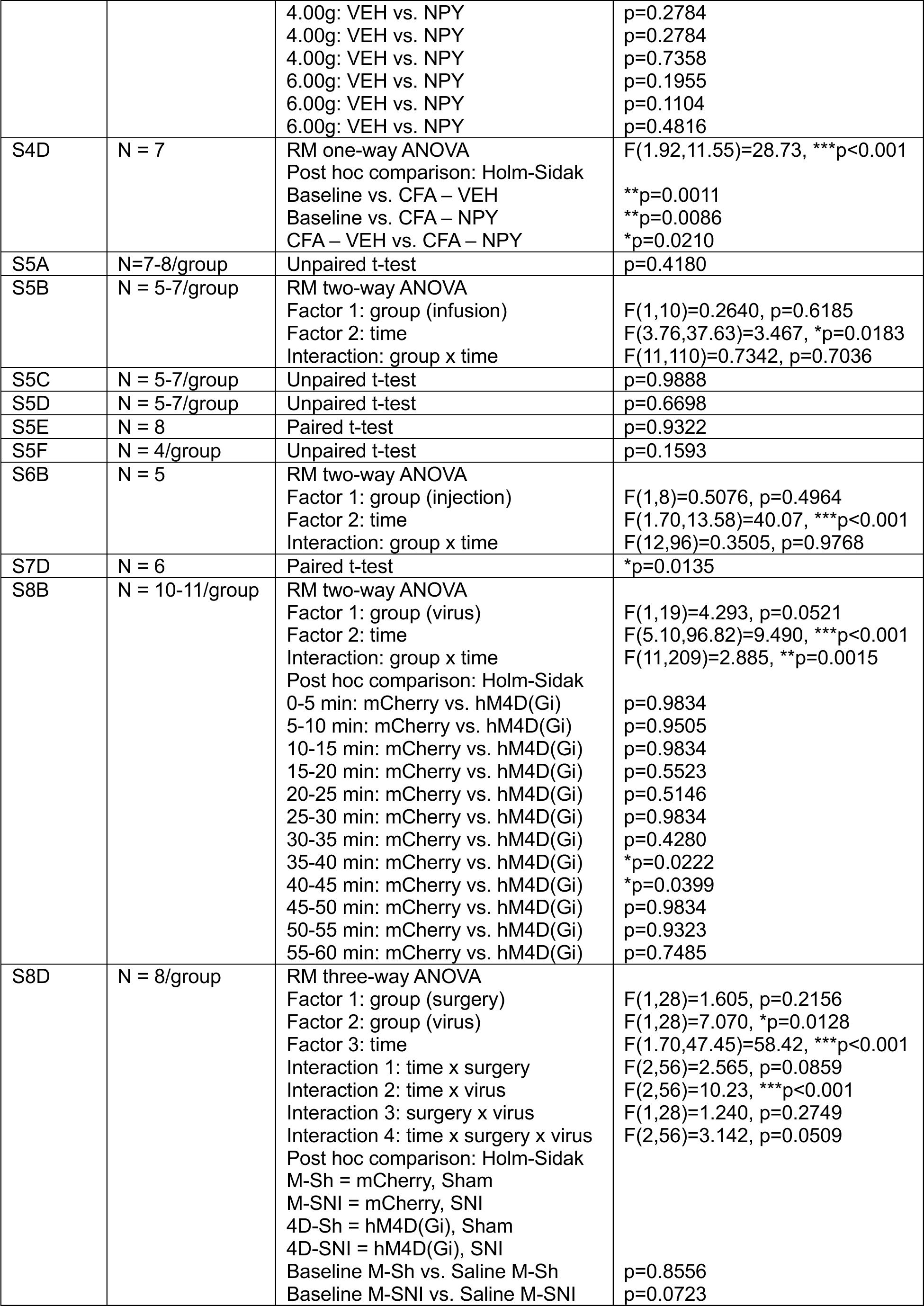
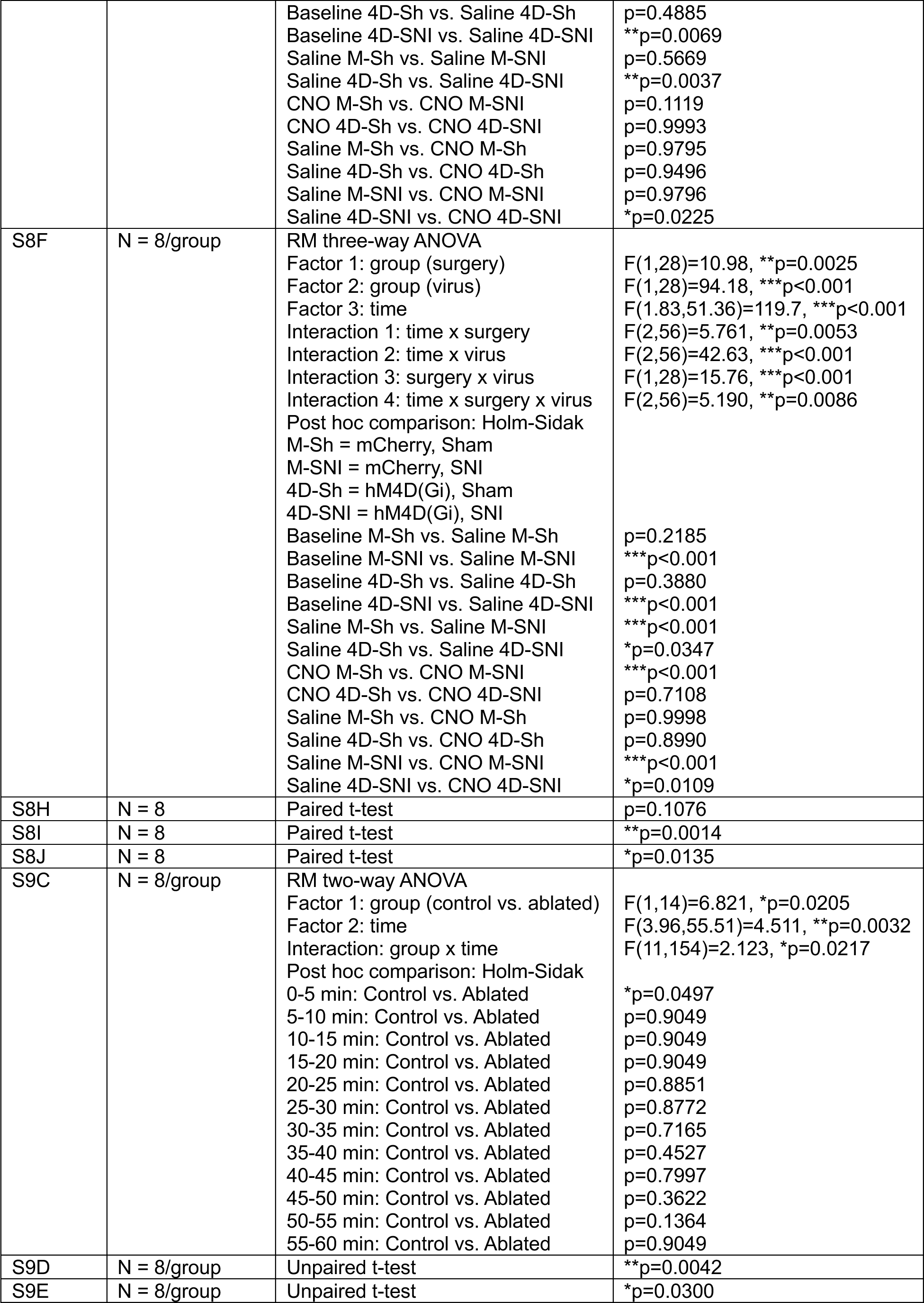
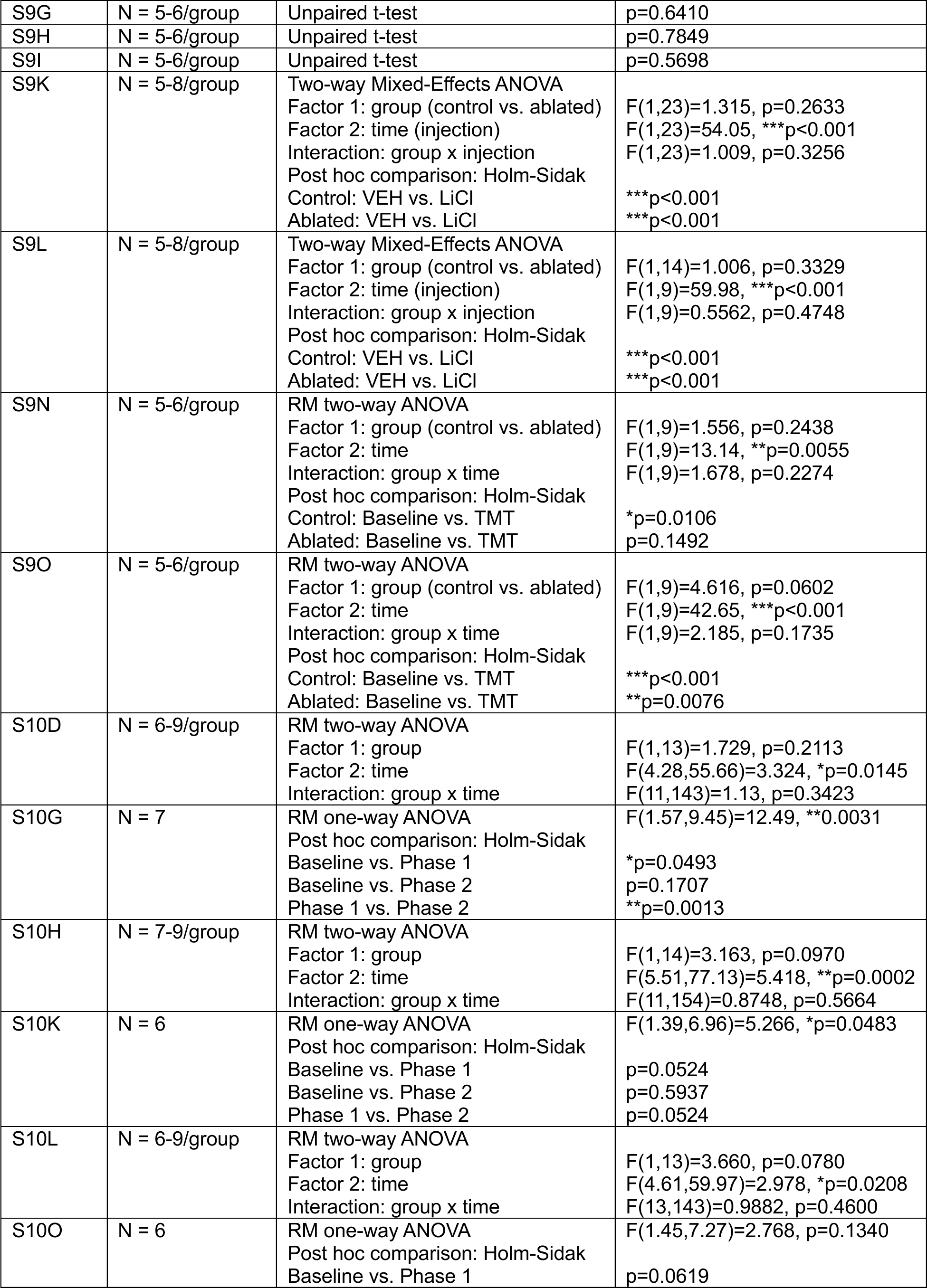
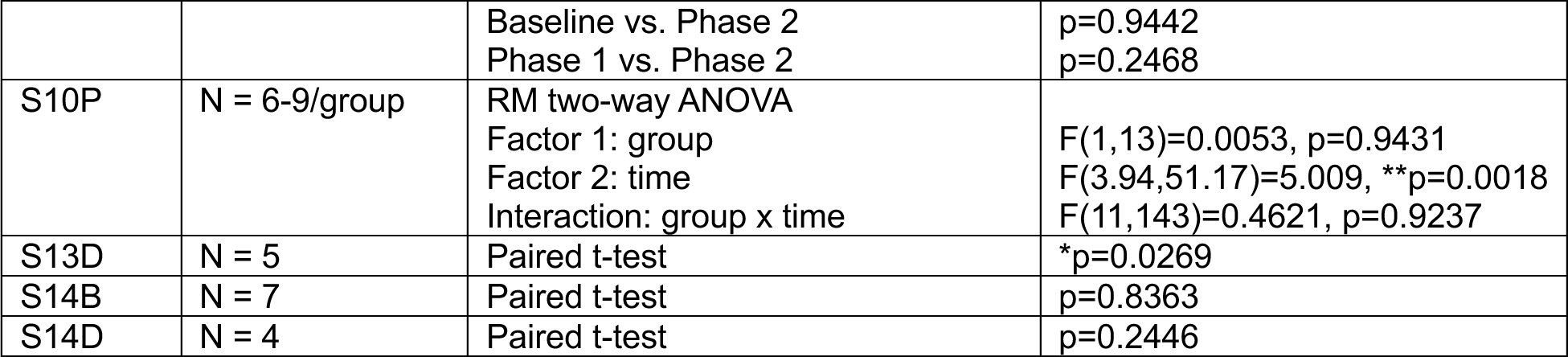
Sample sizes, statistical tests, and results used in supplementary figures.

